# Hyperinsulinemia acts via acinar insulin receptors to initiate pancreatic cancer by increasing digestive enzyme production and inflammation

**DOI:** 10.1101/2022.05.06.490845

**Authors:** Anni M.Y. Zhang, Yi Han Xia, Jeffrey S.H. Lin, Ken H. Chu, Wei Chuan K. Wang, Titine J.J. Ruiter, Jenny C.C. Yang, Nan Chen, Justin Chhuor, Shilpa Patil, Haoning Howard Cen, Elizabeth J. Rideout, Vincent R. Richard, David F. Schaeffer, Rene P. Zahedi, Christoph H. Borchers, James D. Johnson, Janel L. Kopp

**Affiliations:** Department of Cellular and Physiological Sciences, Life Sciences Institute, University of British Columbia, Vancouver, Canada V6T 1Z3; Department of Pathology and Laboratory Medicine, University of British Columbia, Vancouver, Canada V6T 1Z7; Segal Cancer Proteomics Centre, Lady Davis Institute, Jewish General Hospital, McGill University, Montreal, Canada H3T 1E2; Department of Internal Medicine, University of Manitoba, Winnipeg, Canada R3A 1R9; Manitoba Centre for Proteomics and Systems Biology, Winnipeg, Canada R3E 3P4; Gerald Bronfman Department of Oncology, Jewish General Hospital, McGill University, Montreal, Canada H4A 3T2

**Keywords:** pancreatic ductal adenocarcinoma, PanIN, hyperinsulinemia, obesity, insulin resistance

## Abstract

The rising pancreatic cancer incidence due to obesity and type 2 diabetes is closely tied to hyperinsulinemia, an independent cancer risk factor. Previous studies demonstrated reducing insulin production suppressed pancreatic intraepithelial neoplasia (PanIN) pre-cancerous lesions in *Kras*-mutant mice. However, the pathophysiological and molecular mechanisms remained unknown and in particular it was unclear whether hyperinsulinemia affected PanIN precursor cells directly or indirectly. Here, we demonstrate that insulin receptors (*Insr*) in *Kras*^G12D^-expressing pancreatic acinar cells are dispensable for glucose homeostasis, but necessary for hyperinsulinemia-driven PanIN formation in the context of diet-induced hyperinsulinemia and obesity. Mechanistically, this was attributed to amplified digestive enzyme protein translation, triggering of local inflammation and PanIN metaplasia *in vivo*. *In vitro*, insulin dose-dependently increased acinar-to-ductal metaplasia formation in a trypsin– and *Insr*-dependent manner. Collectively, our data shed light on the mechanisms connecting obesity-driven hyperinsulinemia and pancreatic cancer development.

## INTRODUCTION

The 5-year survival rate of pancreatic ductal adenocarcinoma (PDAC) is less than 10% and it is projected to become the 2^nd^ leading cause of cancer death by 2030^1^. Chronic pancreatitis, family history, smoking, obesity, and Type 2 Diabetes (T2D) are risk factors for pancreatic cancer^2^. Obesity and T2D are usually accompanied by hyperinsulinemia, hyperglycemia, increased inflammation, and dyslipidemia, which have all been proposed as underlying factors that drive the increased PDAC morbidity and mortality in this patient population^3,4^. Epidemiological studies consistently show that hyperinsulinemia is associated with increased risk of developing PDAC and worse survival^5,6^. Consistent with these epidemiological studies, experimentally inducing a hyperinsulinemic state in mice by feeding them a diet high in fat content consistently promotes tumor initiation in the pancreas when compared to chow fed mice^7,8^. Furthermore, we previously demonstrated that genetically limiting the high-fat diet mediated increases in insulin, reduced PDAC development, independently of hyperglycemia^9,10^. Single-cell analysis revealed that hyperinsulinemia altered gene expression in multiple cell types in the PanIN microenvironment^10^, leaving open the question of whether the protective effects of reduced insulin production are direct on the tumor precursor cells or whether they are mediated indirectly by local immune cells, local fibroblasts, and/or via distant effects on adiposity^11–13^. Consistent with a direct effect on the epithelium, insulin stimulates proliferation in the PANC-1 and HPDE cell lines *in vitro*^14^, but this does not provide information on the initiation of PDAC *in vivo*.

Altered insulin/IGF signaling, which includes cascades involving KRAS/MAPK/ERK and PI3K/AKT/mTOR, is prominent in human and animal pancreatic cancer. Activating mutations in *KRAS* are detected in >95% of PDAC clinical cases and induce PanIN pre-cancerous lesions and tumours in mice^15^. Activating mutations in *PIK3CA* are found in 3-5% of PDAC patients^16,17^ and can initiate PDAC in mice^18^, while silencing *Pik3ca* was protective in *Pdx1*-Cre;*Kras*^LSL-G12D^;*Trp53*^LSL-R172H^ mice^19^. Strategies that systemically reduce signaling downstream of Insr/Igf1r can suppress PDAC^18,20^, but they do not distinguish between the roles for insulin, IGFs, or other upstream growth factors. Despite indirect evidence for an important role of insulin/IGF signaling in this and other cancers^3,4^, a direct and causal role for the insulin receptor (*Insr*) alone has not been demonstrated for any cancer. Insr protein is increased in some breast, prostate, and liver cancers^21–23^, but its role in the pancreas remains enigmatic. In this study, we tested the hypothesis that hyperinsulinemia-induced enhancement of PDAC initiation is mediated through direct Insr signaling in pancreatic acinar cells. We found mice that consumed a high fat diet had a significant reduction in PanIN and tumor development when they lacked *Insr* specifically in Kras^G12D^-expressing acinar cells. These findings indicate that hyperinsulinemia directly contributes to pancreatic cancer initiation through Insr in acinar cells via a mechanism that involves increased production of digestive enzymes and subsequent pancreatic inflammation.

## RESULTS

### Effects of acinar specific *Insr* loss on body weight and glucose homeostasis

To test our primary hypothesis, that hyperinsulinemia drives pancreatic cancer development via Insr cell autonomously, we generated mouse models in which *Kras*^G12D^ expression^24^ and loss of *Insr* were both induced in acinar cells. Our cohorts contained mice with full *Insr* gene dosage, *Ptf1a*^CreER^;*Kras*^LSL-G12D^;*Insr*^w/w^;nTnG (PK-*Insr*^w/w^); mice with partially reduced *Insr*, *Ptf1a*^CreER^;*Kras*^LSL-G12D^;*Insr*^w/f^;nTnG (PK-*Insr*^w/f^); or knockout mice without any *Insr*, *Ptf1a*^CreER^;*Kras*^LSL-G12D^;*Insr*^f/f^;nTnG (PK-*Insr*^f/f^) in Kras^G12D^-expressing acinar cells (Figure 1A). We also generated a limited number of *Ptf1a*^CreER^;*Insr*^w/w^;nTnG and *Ptf1a*^CreER^;*Insr*^f/f^;nTnG mice to assess the baseline roles of *Insr* in acinar cells (Figure 1A). All mice were fed a high fat diet (HFD) (Figure 1B), which is known to produce sustained hyperinsulinemia and accelerate PanIN/PDAC development^7,8^. Tamoxifen was injected at 4 weeks of age to simultaneously induce expression of mutant *Kras*^G12D^ and nuclear GFP^25^, as well as deletion of floxed *Insr* alleles, specifically in acinar cells (Figure 1B). As expected, mice consuming HFD gained weight over time (Figure 1C-D) and at 43.5 weeks weighed more than the average 38– or 33-gram weight expected for chow-fed 44-week-old male or female C57Bl6J mice, respectively^26^ (see also data at Jax.org). Acinar *Insr* loss did not significantly affect the weight gain of male or female mice in the context of mutant *Kras* (Figures 1C-D), but the presence of mutant *Kras* allele affected the weight gain of males over time compared to *Kras* wildtype mice (Figure 1C). This is consistent with previous reports that acinar-to-ductal metaplasia (ADM)– and PanIN-mediated disruption of pancreatic function blunted weight gain in *Pdx1Cre;Kras*^G12D^ mice^27^. Fasting glucose was not different between genotypes (Figures 1E-F). HFD induced similar levels of hyperinsulinemia in both genotypes (Figures 1G-H). As expected^9,10^, males had higher insulin levels compared to females (Figures 1G-H). These insulin levels were well above chow diet-fed male C57Bl6J mice (138 +/− 24 pM)^28^. Our data demonstrate that acinar-specific *Insr* deletion does not have major effects on glucose homeostasis in mice. Thus, our model enabled us to test the role of acinar cell *Insr* in the context of hyperinsulinemia and normoglycemia, modelling the typical obese, non-diabetic state.

**Fig. 1.**
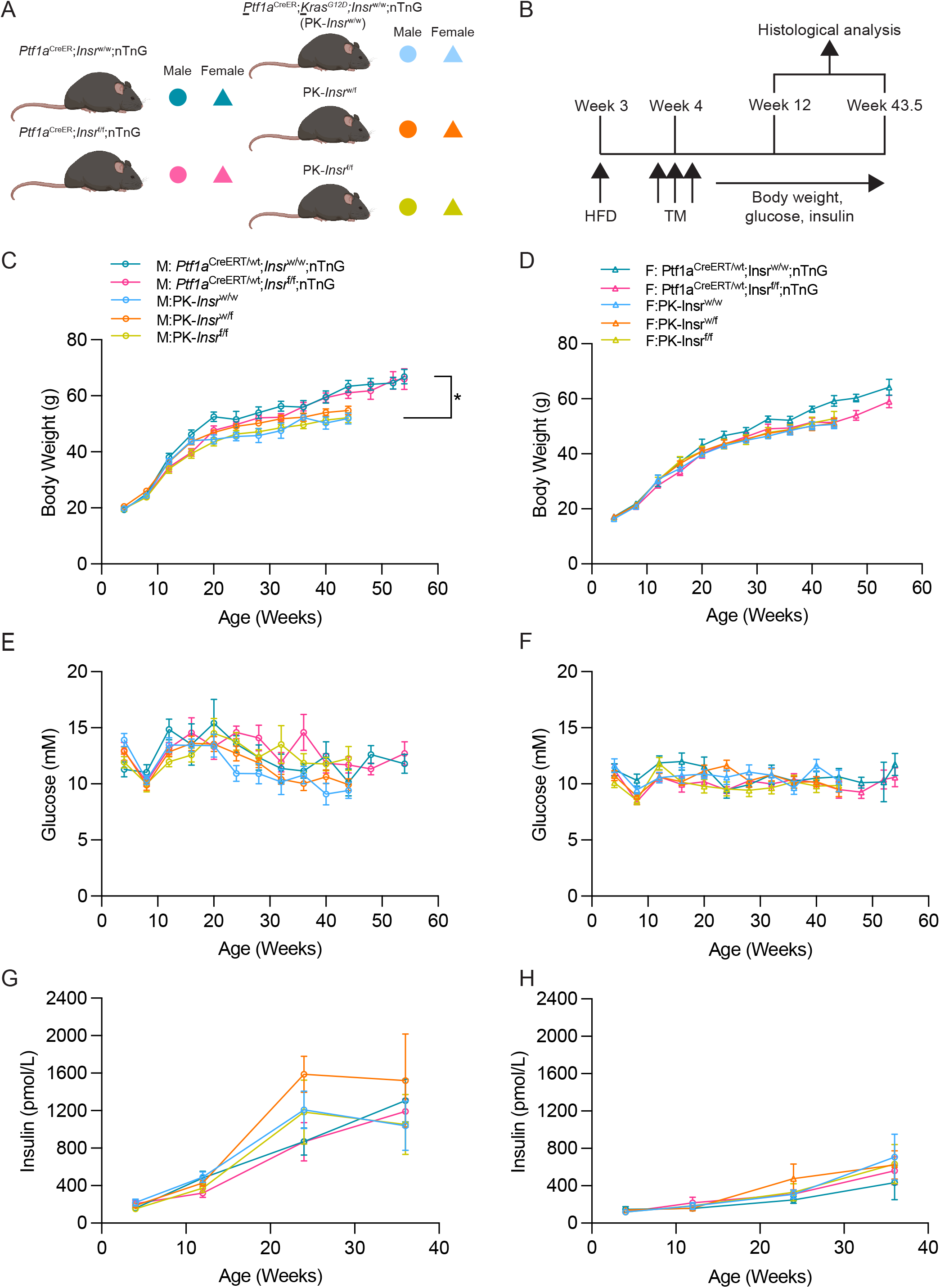
Loss of *Insr* in pancreatic acinar cells had no effect on glucose homeostasis and pancreas size. **A**, Schematic describing a mouse model designed to test the role of insulin receptor signaling on HFD-accelerated PDAC initiation. On the background of the *Ptf1a*^CreER^ mice, we generated mice having two wild-type *Insr* alleles (*Ptf1a*^CreER^;*Insr*^w/w^;nTnG) or two *Insr* floxed alleles (*Ptf1a*^CreER^;*Insr*^f/f^;nTnG). On the background of the *Ptf1a*^CreER^-induced *Kras*^G12D^ pancreatic cancer model with the nTnG lineage allele (PK), we generated mice having two wild-type *Insr* alleles (PK-*Insr*^w/w^), one *Insr* floxed allele (PK-*Insr*^w/f^), or two *Insr* floxed alleles (PK-*Insr*^f/f^). Both male (M:, circles) and (F:, triangles) mice were analyzed. **B**, Three-week-old mice were weaned and provided high fat diet (HFD) for the duration of the study. At 4-weeks-old, they were injected with tamoxifen (TM) on 3 consecutive days. Physiological measures were taken every 3 months for 10 months, mice were euthanized at 12 or 43.5 weeks of age for histopathology. The mice which were euthanized at 12 weeks of age were also used for proteomics, phospho-proteomics and RNA-sequencing analyses. **C-H,** Body weight (**C-D**), fasting blood glucose (**E-F**), and fasting insulin (**G-H**) measurements in male (**C, E, G**) and female (**D, F, H**) *Ptf1a*^CreER^-*Insr*^w/w^;nTnG and *Ptf1a*^CreER^-*Insr*^f/f^;nTnG mice measured over 57 weeks (n= 10-17) or PK-*Insr*^w/w^, PK-*Insr*^w/f^, and PK-*Insr*^f/f^ mice up to 43.5 weeks (n= 13-33). There were significant differences between PK-*Insr*^w/w^ and PK-*Insr*^f/f^ compared to *Ptf1a*^CreER^-*Insr*^w/w^;nTnG for male body weight gain over time (p < 0.05). Values shown as mean ± SEM. *p<0.05 by compare groups of growth curves permutation test (C-D) or by mixed effects (E-H). See also Figure S1 and S2.

### Acinar *Insr* knockout limits loss of normal pancreatic parenchyma by Kras^G12D^

To test whether *Insr* loss affected Kras^G12D^-mediated PDAC formation from acinar cells, we examined a cohort of 5-16 PK-*Insr*^w/w^, PK-*Insr*^w/f^, and PK-*Insr*^f/f^ mice of each sex and genotype. We planned to quantify the extent of lesions at ∼1 year of age based on the timing of our previous genetic models with reduced insulin^9,10^, but half the male PK-*Insr*^w/w^ mice and a few female PK-*Insr*^w/w^ mice reached humane endpoint earlier (Figure 2A). Therefore, we chose to end the study at 43.5 weeks of age for Kras^G12D^ mice. Through necropsy and histology, we noted that PDAC was present in 4 male PK-*Insr*^w/w^ mice, but more importantly, almost the entire parenchyma of male PK-*Insr*^w/w^ mice was replaced by ductal metaplasia comprised of cysts, PanIN, and ADM (Figure 2A-C, E). Three male PK-*Insr*^w/f^ mice reached humane endpoint prior to 43.5 weeks without macroscopic tumors (Figure 2A-B), but more animals of this genotype retained normal acinar cell area (Figure 2C, E). No male PK-*Insr*^f/f^ mice reached humane endpoint by 43.5 weeks of age (Figure 2A-B) and they retained even more normal acinar cell area on average (Figure 2C, E). One female PK-*Insr*^w/w^ mouse, as well as a 1 female PK-*Insr*^w/f^ mouse, reached humane endpoint prior to 43.5 weeks of age, while no female PK-*Insr*^f/f^ mice reached humane endpoint by 43.5 weeks of age (Figure 2A-B). No macroscopic tumors were noted at necropsy (Figure 2A-B), but histologically, we found that the incidence of PDAC in females was *Insr* dosage-dependent (Figures 2B, and D). The majority of female mice from every genotype retained some normal parenchyma (Figures 2D-E), potentially explaining why females reached humane endpoint less often than males. This is consistent with previous reports suggesting that the timing and/or extent of lesion formation differs between male and female mice in the context of HFD^8^. Together, these data strongly suggested that limiting or eliminating insulin/Insr activity specifically in acinar cells reduced the propensity of HFD and Kras activation to transform the pancreas.

**Fig. 2.**
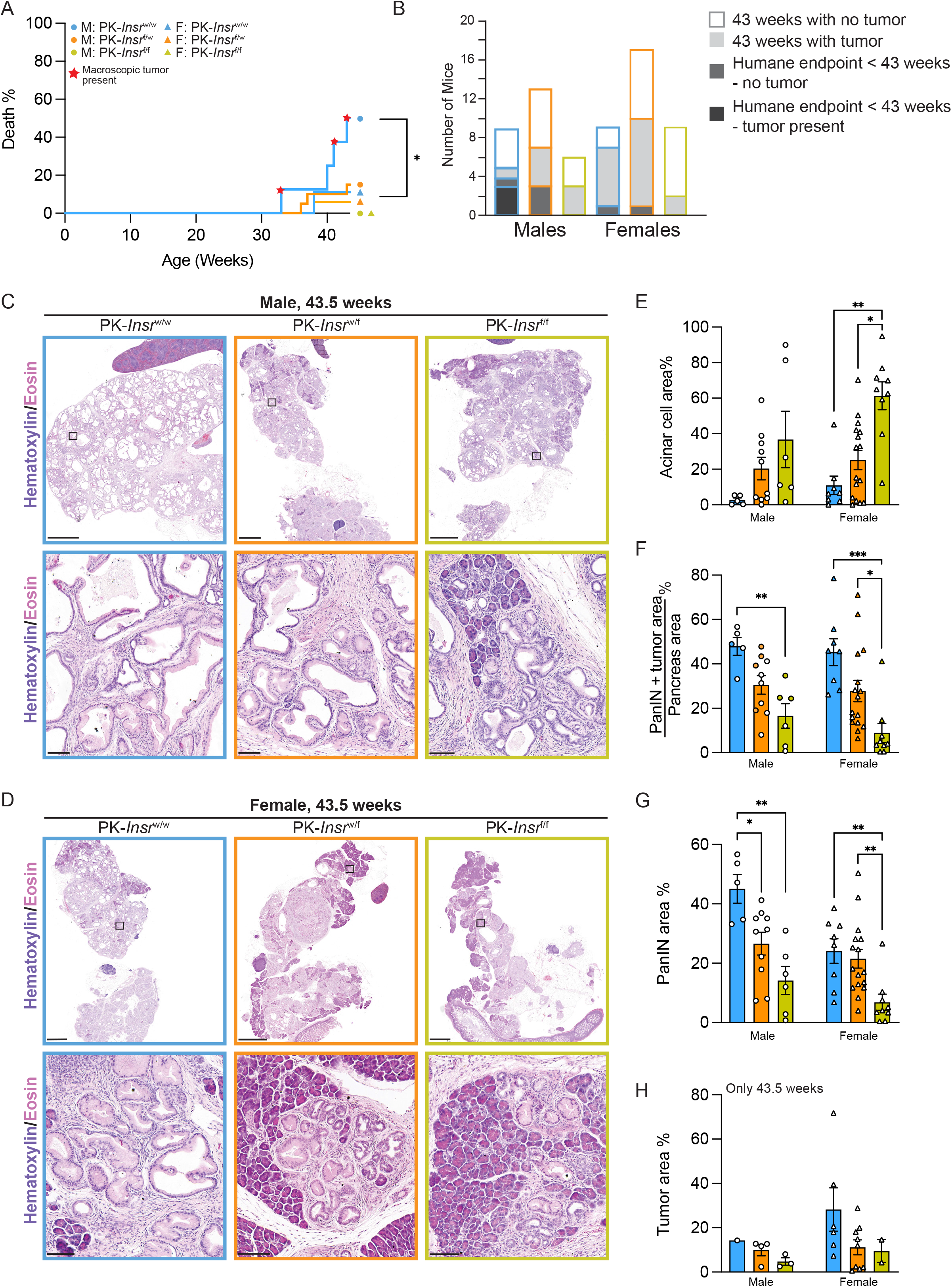
Loss of *Insr* in acinar cells reduced PanIN and PDAC formation. **A**, Percentage of mice of the indicated genotype that were found dead or reached humane endpoint throughout the study. The star symbol indicates whether a macroscopic mass was observed in the pancreas at necropsy, when possible. There were 9 PK-*Insr*^w/w^, 13 PK-*Insr*^w/f^, and 6 PK-*Insr*^f/f^ male mice and 9 PK-*Insr*^w/w^ 17 PK-*Insr*^w/f^, and 9 PK-*Insr*^f/f^ female mice in the cohort. A significant difference was detected between the curves and only the male PK-*Insr*^w/w^ mice obtained a media survival value. At 43.5 weeks, the remaining mice were euthanized and assessed for presence of tumors. **B,** The number of mice dead before or at 43 weeks with or without tumor presence. **C-D,** Representative whole section (top) and high-magnification (bottom) H&E images of pancreatic slides from 43.5-week-old male (**C**) and female (**D**) PK-*Insr*^w/w^, PK-*Insr*^w/f^, and PK-*Insr*^f/f^ mice. **E-H,** Quantification of acinar cell area (**E**), PanIN plus tumor area (**F**), PanIN only area (**G**), or tumor area (**H**) in pancreata from each genotype and sex (male or female) at 43.5 weeks of age (n= 5-16). (**F**) Filled circles or triangles denote mice that developed tumors. Scale bars: 2 mm (**C-D,** top) and 0.1mm (**C-D**, bottom). Values are shown as mean ± SEM. *p<0.05, **p<0.01, ***p<0.001 by Log-rank (Mantel-Cox) test (A), or by Kruskal-Wallis test (E-G). See also Figure S3.

### Loss of *Insr* in acinar cells reduces the extent of *Kras*^G12D^-mediated lesion formation

We next quantified the extent of the tissue disruption and lesion formation only in mice that were 43.5 weeks of age to assess the extent of disease in the presence or absence of *Insr*. In parallel with PanIN quantification, we stained all pancreata for nuclear GFP from the nTnG lineage reporter allele^25^. As expected, most acinar cells and PanIN lesions were GFP positive (Figures S1A-B), confirming that the lesions arose from Ptf1a^+^ acinar cells. Pancreata from *Kras* wild-type *Ptf1a*^CreER^;*Insr*^w/w^;nTnG and *Ptf1a*^CreER^;*Insr*^f/f^;nTnG mice were similarly comprised of acinar cells and endocrine islets, and had broad GFP expression in acinar cells (Figures S1C-F). In contrast, pancreata from male and female PK-*Insr*^w/w^, PK-*Insr*^w/f^, and PK-*Insr*^f/f^ mice all contained ductal lesions with histological characteristics of metaplastic ducts, including ADM, low-grade and high-grade PanIN, and sometimes PDAC tumors (see above; Figures 2C-D). Reducing *Insr* in Kras^G12D^-expressing acinar cells decreased the area of PanINs plus tumors, PanIN alone, or tumor area in a dose dependent manner in males and females (Figures 2F-H). Consistent with this histological-based quantification, measuring the pancreatic area containing ductal metaplasia (ducts, ADM, PanIN and PDAC) or mucinous lesions (PanIN and some tumors) with Ck19 or Alcian blue staining, respectively, similarly showed that female PK-*Insr*^w/w^ mice formed significantly more lesions than female PK-*Insr*^f/f^ mice (Figures S2A-F). Notably, male PK-*Insr*^w/w^ mice had a significantly higher amount of Ck19^+^ area, but a similar amount of Alcian blue^+^ area, compared to other genotypes (Figure S2A-F). This latter observation is likely explained by the presence of Alcian blue negative high-grade PanIN lesions and large cysts with predominantly normal ductal epithelium, which were more prevalent in male than female PK-*Insr*^w/w^ pancreata (Figures 2C-D, S2A-F). Indeed, most male PK-*Insr*^w/w^ and some female PK-*Insr*^w/w^ pancreata were comprised of almost all Ck19^+^ area with little acinar cell area left, while larger areas of normal acinar cells correlated with lower Ck19^+^ areas in PK-*Insr*^w/f^ and PK-*Insr*^f/f^ mice (Figures 2C-E, S2G-H). Therefore, our data strongly suggested that acinar cell *Insr* dose-dependently regulates oncogenic Kras-induced PanIN formation in the context of diet-induced hyperinsulinemia.

### Loss of *Insr* in *Kras*^G12D^-expressing acinar cells blocks early PanIN initiation

To examine whether early PanIN initiation was specifically affected by loss of *Insr* in acinar cells, we examined mice at 12 weeks of age. At 8 weeks post-tamoxifen injection, acinar cells and PanIN lesions were also GFP positive indicating good recombination efficiency (Figure S3A-B). There were no significant differences in pancreas weight or Cpa1^+^ area between PK-*Insr*^w/w^, PK-*Insr*^w/f^, and PK-*Insr*^f/f^ mice in either sex, nor were there any differences in pancreatic weight of Kras^G12D^-expressing mice compared to *Ptf1a*^CreER^;*Insr*^w/w^;nTnG or *Ptf1a*^CreER^;*Insr*^f/f^;nTnG mice (Figure S3C-G). This is consistent with previous data showing that the young pancreata from *Ptf1a*^CreER^;*Kras*^LSL-G12D^ mice have relatively fewer PanIN lesions^24^. Pancreata from PK-*Insr*^f/f^ mice were predominantly normal and had significantly fewer ductal lesions with ADM or low-grade PanIN characteristics when compared to PK-*Insr*^w/w^ mice (Figure 3A-C). Consistent with the increased number of lesions in PK-*Insr*^w/w^ compared to PK-*Insr*^f/f^ mice, there was slightly less Cpa1^+^ area in female PK-*Insr*^w/w^ compared to PK-*Insr*^f/f^ pancreata (Figure S3C-E) and a weak inverse correlation of overall PanIN area to Cpa1^+^ area only for PK-*Insr*^w/w^ mice (Figure S3F). Bulk RNA sequencing and gene ontology analyses also showed that PK-*Insr*^w/w^ compared to PK-*Insr*^f/f^ pancreata had higher expression of genes associated with Ras signaling, proliferation, and inflammation, which are typically associated with PanIN initiation^29^ (Figure 3D-E, Table S1). The significant inhibition of PanIN formation at 12 weeks of age solely through reducing *Insr* in acinar cells strongly supported a model in which diet-induced hyperinsulinemia promoted tumor initiation through insulin receptor action on acinar cells.

**Fig. 3.**
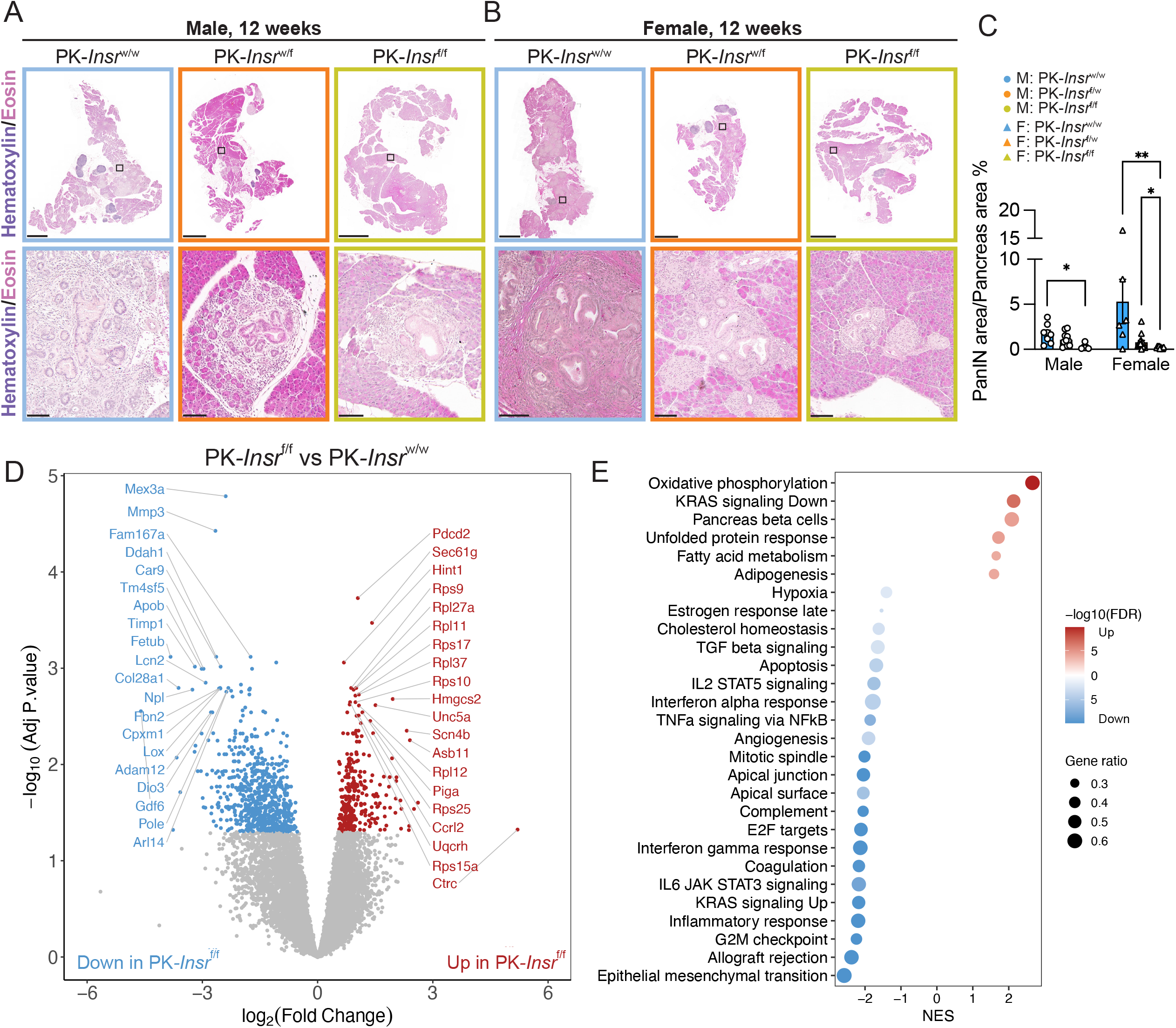
Loss of *Insr* in acinar cells reduced PanIN lesions initiation. **A-B**, Representative whole section (top) and high-magnification (bottom) H&E images of pancreatic slides from 12-week-old male (**A**) and female (**B**) PK-*Insr*^w/w^, PK-*Insr*^w/f^, and PK-*Insr*^f/f^ mice. **C,** Quantification of PanIN area in 12-week-old pancreata from each genotype and sex (male or female) (n= 3-14). The difference in PanIN formation between male and female PK-*Insr*^w/w^ mice was not significantly different. **D**, Volcano plot for genes that were significantly up– or down-regulated in PK-*Insr*^f/f^ mice (n=4) compared to PK-*Insr*^w/w^ mice (n=4). **E,** Gene set enrichment analysis for genes differentially expressed between PK-*Insr*^f/f^ mice and PK-*Insr*^w/w^ mice. Scale bars: 2 mm (top) and 0.1mm (bottom). Values are shown as mean ± SEM. *p<0.05, **p<0.01 by Kruskal-Wallis test (C). See also Figure S4 and Table S1.

### Akt and Erk signaling are reduced in PanINs in the absence of *Insr*

Insulin receptor tyrosine kinases act through the PI3K/Akt/mTor and Mapk/Erk signaling cascades. To further assess the molecular changes associated with the loss of *Insr* in acinar cells, we examined the phosphorylation of Akt at S473 and the phosphorylation of Erk at Thr202/Tyr204 by immunohistochemistry (Figure S4). As previously shown^29^, signal for pAkt and pErk was present in PanIN lesions in PK-*Insr*^w/w^ pancreata (Figure S4A-D). As expected in the context of reduced insulin signaling and reduced PanIN area, the pancreatic area positive for both pAkt or pErk signal (Figure S4E-F) was reduced in PK-*Insr*^f/f^ compared to PK-*Insr*^w/w^ pancreata (Figure S4A-D). Although the pAkt and pErk signal also typically decreased or changed localization in remaining PanIN lesions in PK-*Insr*^f/f^ mice, the pAkt^+^ signal was present in the stromal cells surrounding the PanIN lesions regardless of genotype or sex (Figure S4A-D). The variability in levels of pAKT and pERK signaling could be due to incomplete recombination of the *Insr* alleles in the few PanINs that do form in the PK-*Insr*^f/f^ mice, or alternatively other signaling pathways could influence the levels of pAKT and pERK, such as EGFR signaling. These data suggest that loss of *Insr* in acinar cells can reduce PI3K/Akt/mTor and Mapk/Erk signaling during oncogenic Kras^G12D^-mediated PanIN formation.

### Pancreata from mice lacking acinar cell Insr have reduced digestive enzymes

To define the underlying molecular mechanisms of *Insr* action during PanIN initiation, we conducted unbiased total proteomic and phospho-proteomic analyses using the head of the pancreas from the 12-week-old female mice (Figure 3, Tables S2-S3) with and without Kras^G12D^ and/or *Insr*. For the total proteome dataset, we obtained reliable, quantitative data on 2889 proteins across all the samples. First, we examined the effects of mutant Kras in the context of normal *Insr*. Consistent with our histological analyses identifying a change in PanIN frequency, we found that the inflammation– and PanIN-associated proteins, Reg3a, Reg3b, Reg2, Tff1, Gkn1, Gkn2, and Agr2^30–33^, were among the 124 proteins significantly enriched in PK-*Insr*^w/w^ compared to *Ptf1a*^CreER^;*Insr*^w/w^;nTnG pancreata (Figure 4A, Table S2). We also found that proteins associated with acinar cell function, such as Ctrc, Dbi, Pla2g1b, and Clps, were among the 122 proteins significantly down-regulated as a consequence of *Kras*^G12D^ expression in acinar cells (Figure 4A, Table S2). Other proteins associated with acinar cell function, like Cpa1, Spink1, and Cela1, remained unchanged between the PK-*Insr*^w/w^ and *Ptf1a*^CreER^;*Insr*^w/w^;nTnG genotypes (Table S2). We also compared the differentially expressed pathways between PK-*Insr*^w/w^ and *Ptf1a*^CreER^;*Insr*^w/w^;nTnG mice to differentially expressed pathways between human PDAC and normal pancreas tissues adjacent to human PDAC^34^ and found concordance (Figure S5A-B), reinforcing the translational relevance of this model. Interestingly, we also found 92 proteins increased in abundance and 155 proteins had decreased abundance solely due to loss of *Insr* in Kras wild-type acinar cells (Figure 4B), with the caveat that we analyzed a low sample number. However, reductions in Ctrc, Clps, Pla2g1b in this analysis (Figure 4B, Table S2) suggested that *Insr* may have a role in regulating the function of wild-type acinar cells in mice fed HFD.

**Fig. 4.**
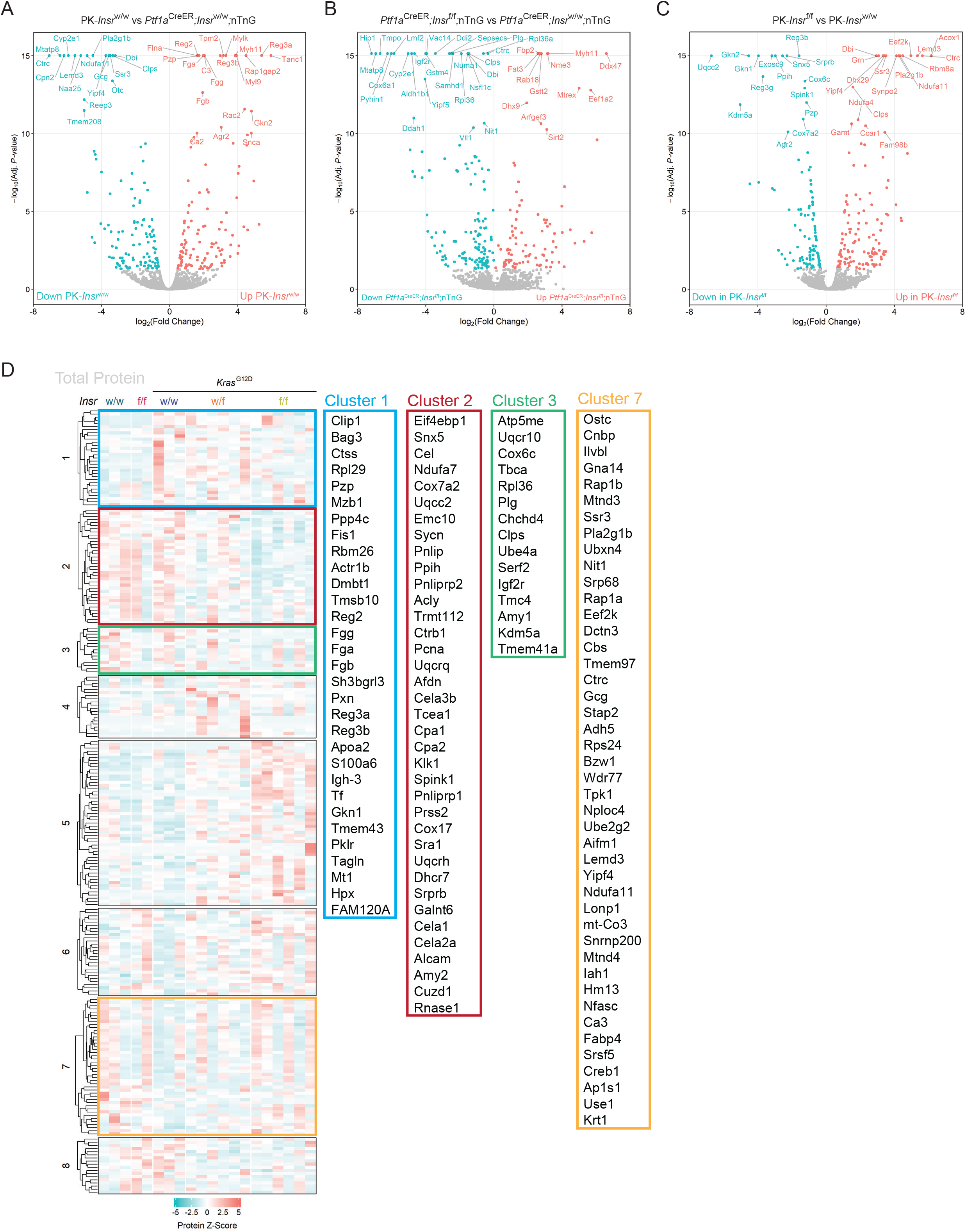
The proteomic analysis of insulin receptors’ effects on PanIN initiation. **A**, Volcano plot for proteins that were significantly up– or down-regulated in PK-*Insr*^w/w^ mice (n=3) compared to *Ptf1a*^CreER^;*Insr*^w/w^;nTnG mice (n=3). **B**, Volcano plot for proteins that were significantly up– or down-regulated in *Ptf1a*^CreER^;*Insr*^f/f^;nTnG mice (n=2) compared to *Ptf1a*^CreER^;*Insr*^w/w^;nTnG mice (n=3). **C**, Volcano plot showing proteins that were significantly up– or down-regulated in PK-*Insr*^f/f^ mice (n=6) compared to PK-*Insr*^w/w^ mice (n=3). **D,** Proteins that were differentially abundant between PK-*Insr*^w/w^ and PK-*Insr*^f/f^ mice were selected and their average abundance across all genotypes was used for k-means clustering. The clusters were visualized by heatmap to show their variance across all genotypes and samples. The proteins in clusters 1, 2, 3, and 7 are listed in the order shown in the heatmap from top to bottom. This information can also be found in the Supplemental Table S4 Clusters 1-8. See also Figure S5 and Tables S2.

To define the molecular mechanisms associated with *Insr* deletion in the context of mutant Kras, we focused further analyses on comparing the PK-*Insr*^f/f^ pancreata to the PK-*Insr*^w/w^ pancreata. We found that 135 proteins were enriched and 117 were depleted in PK-*Insr*^f/f^ mice compared to PK-*Insr*^w/w^ controls (Figure 4C). We then performed a K-means clustering analysis with the abundance values of these differentially enriched proteins in all genotypes to identify groups of proteins that varied by *Insr* status, Kras mutation, or both (Figure 4D and Table S4-all clusters). In order to investigate potential functional enrichment of protein groups based on cell signaling, intracellular localization, and biological process, we also used the differentially abundant proteins between PK-*Insr*^w/w^ and PK-*Insr*^f/f^ mice to build protein-protein interaction networks using STRING^35^ and assigned these proteins to their intracellular organelle locations in a diagram using the COMPARTMENTS section of GeneCards and/or existing knowledge of their function^36^ (Figure 5A). As expected from our histological analyses, the total proteomics analyses confirmed that proteins associated with PanIN initiation or formation, such as Reg3a, Reg3b, Reg2, Tff1, Gkn1, and Gkn2, were higher in PK-*Insr*^w/w^ pancreata compared to *Ptf1a*^CreER^*;Insr*^w/w^;nTnG and PK-*Insr*^f/f^ pancreata (Figure 4C, 4D-cluster 1, 5A). Proteomic analyses highlighted the striking downregulation of the majority of the proteins packaged into zymogen granules for secretion in PK-*Insr*^f/f^ compared to *Ptf1a*^CreER^;*Insr*^w/w^;nTnG and PK-*Insr*^w/w^ mice (Figure 4D-cluster 2,3, 5A-B). Acinar cells produce a large amount of protein every day and in *Ptf1a*^CreER^;*Insr*^w/w^;nTnG mice and PK-*Insr^w/w^* pancreata fed HFD ∼19% of all the peptides detected in our analyses were associated with the zymogen granules (Figure 5C). However, this percentage was reduced significantly to 12.2% in PK-*Insr*^f/f^ pancreata (Figure 5C). This reduction in zymogen granule proteins is likely an underestimation of the effect on the total proteome, as our differential abundance analyses were normalized to total protein content. In sum, loss of *Insr* in acinar cells resulted in a coordinated decrease in digestive enzymes produced by acinar cells in the context of HFD.

**Fig. 5.**
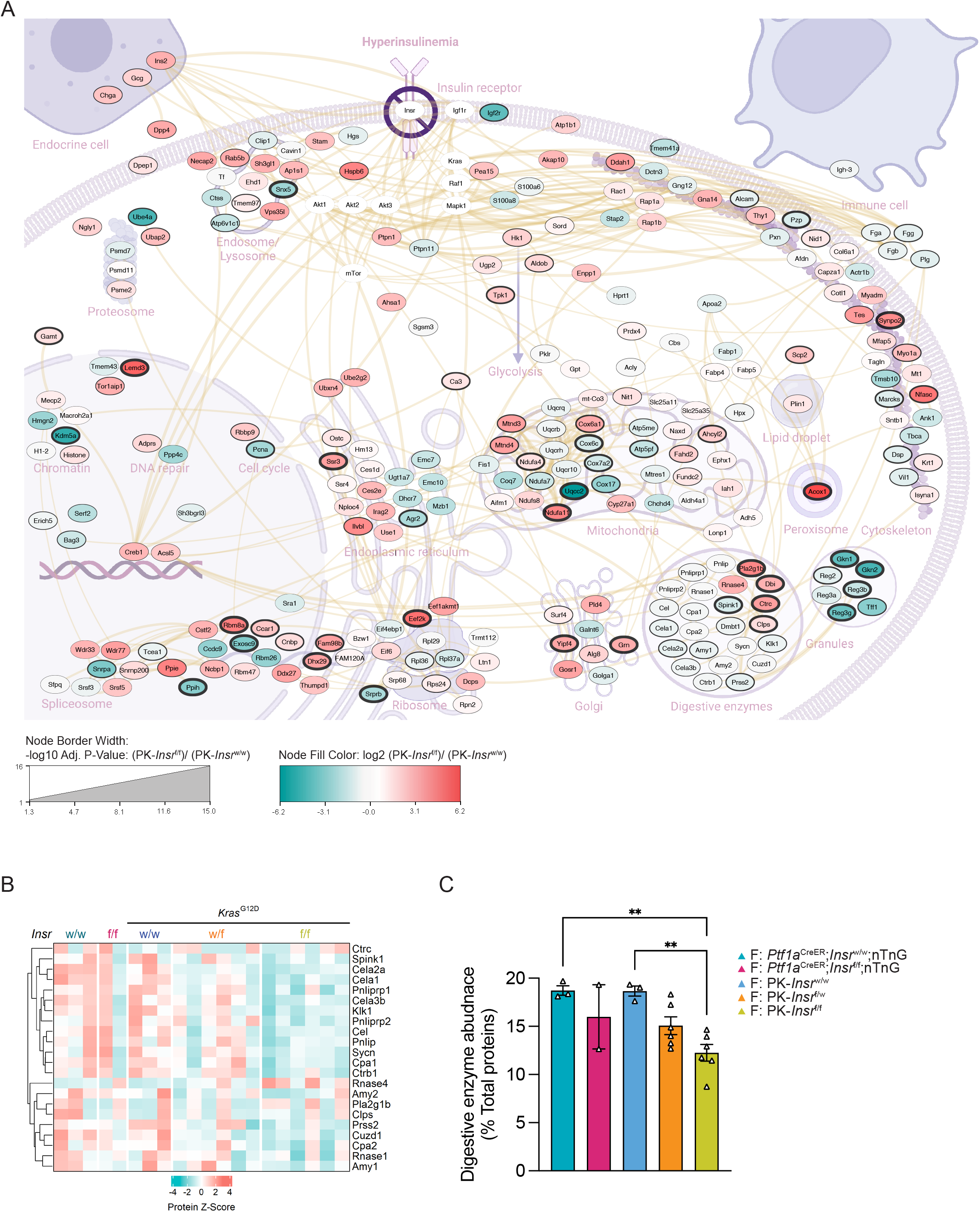
Loss of *Insr* in acinar cells reduced digestive enzymes production in pancreas. **A**, Differentially abundant proteins between PK-*Insr*^w/w^ and PK-*Insr*^f/f^ pancreata were connected using STRING and diagrammatically presented here in the context of Insr signaling and cellular organelles and functions (outline created in Biorender). The color in each oval reflects the fold change, while the thickness of the line around the oval represents the Adj. p-value. Proteins depicted in white ovals were added to show the potential relationship of the differentially abundant proteins to insulin signaling. **B**, Heatmap showing the change of protein abundance between genotypes for proteins involved in pancreatic digestive enzymes secretion. **C**, The percentage of summed digestive enzyme abundances in total measured proteins for each genotype. Values are shown as mean ± SEM. **p<0.01 by one-way ANOVA.

### Insr signaling regulates the post-transcriptional synthesis of digestive enzymes

Pancreatic acinar cells represent the cellular majority in our bulk tissue proteome. With the caveat that our bulk tissue includes multiple cell types, our STRING network analysis nevertheless highlighted mechanisms downstream of *Insr* loss that underlie this dramatic reduction in zymogen granule content in acinar cells. This included increases in proteins with key roles in suppressing protein synthesis at the ribosome (eEF2K), as well as increases in the first enzyme of the fatty acid beta oxidation pathway (Acox1), a protein regulating lysosome function (Grn), and protein processing machinery at the ER and Golgi (Ilvbl, Ssr3, Ssr4, Irag2). Many components of mitochondrial electron transport complexes were changed, as well as endosomal and cytoskeletal proteins, many of which are involved in moving organelles within the cells or in exocytosis, such as Snx5, Vps35l, Sycn (Figure 5A). Finally, we also found decreases in critical parts of the spliceosome (Snrpa), the signal recognition particle complex receptor (Srprb), and components of the large ribosome (Rpl29, Rpl36, Rpl37a). These observations suggested that loss of *Insr* in acinar cells and a reduction in PanIN formation was associated with a post-transcriptional reduction in the synthesis of digestive enzymes.

### *Insr* loss in acinar cells reduced phosphorylation of proteins involved in translation

Parallel phosphoproteomic analysis identified 225 downregulated phospho-peptides and 177 upregulated phospho-peptides by comparing the PK-*Insr*^f/f^ pancreas to PK-*Insr*^w/w^ controls (Figure 6A-D, Table S3). In general, statistically significant phospho-peptide differences were not due to underlying differences in total protein abundance (Figure 6B, Table S5). As above, we mapped the location and function of the modified proteins (Figure 6C-D). Specifically, we found significant decreases in phosphorylation in PK-*Insr*^f/f^ compared PK-*Insr*^w/w^ pancreata for proteins involved in transcription elongation (Eloa, Top2a, Supt5h), mRNA splicing and nuclear speckle formation (Srpk1, many Srsf proteins, Cherp, Srrm1 and Srrm2), as well as protein translation initiation (Eif4b, Eif5b, and eIF3 complex proteins) and elongation (Eef1b, Eef1d, and many ribosomal subunits). There were also decreases in phospho-peptides for Cavin3, which has been implicated in Akt-Erk signaling bias^37^, and the PP1 inhibitor Ppp1r2, at sites known to be influenced by insulin signaling (Figure 6D). There was also reduced phosphorylation of Larp1 at sites known to be regulated by mTorc1 component, Raptor^38^. Larp1 is an RNA-binding protein that links mTorc1 to the translational regulation of terminal oligopyrimidine tract containing mRNAs that encode for ribosomal proteins and elongation factors^39^. Together, our proteomics data suggested that Insulin/Insr promotes the production of proteins in acinar cells in part through its modulation of Larp1 and many other components controlling transcription, translation, and secretion of digestive enzymes. Interestingly, multiple sites on the cholecystokinin (Cck) receptor (Cckar) also showed altered phosphorylation (Figure 6C-D), including sites in the main intracellular loop that are indicative of ligand-induced desensitization^40^ and phospho-sites in the C-terminal tail that have not been previously reported. Collectively, these unbiased and quantitative (phospho)proteomic analyses delineate molecular mechanisms by which hyperinsulinemia, acting via the Insr, may promote Kras-driven pancreatic cancer initiation.

**Fig. 6.**
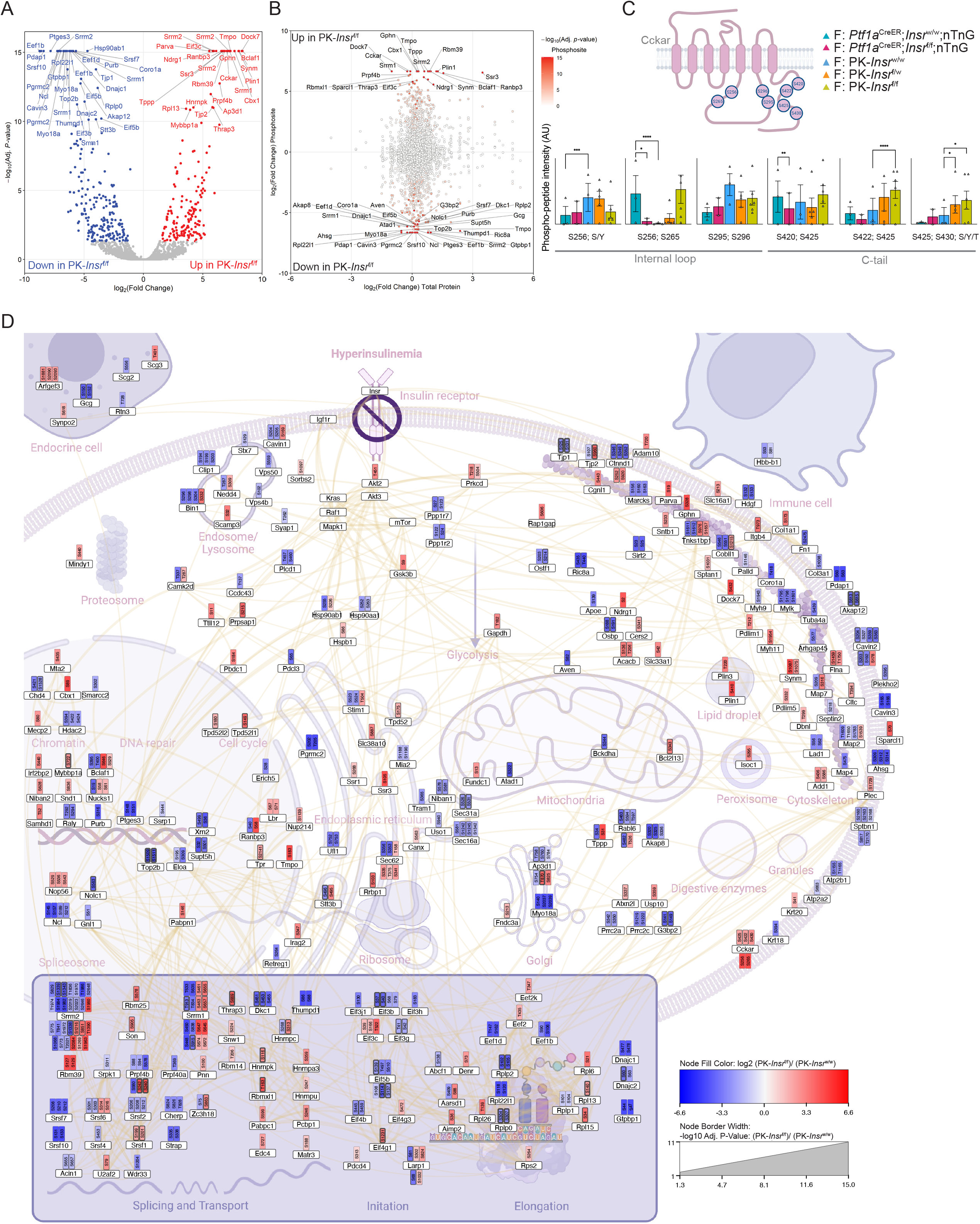
The phospho-proteomic analysis of insulin receptors’ effects on PanIN initiation. **A**, Volcano plots for proteins with phosphor-sites that were significantly up or down regulated in PK-*Insr*^f/f^ mice compared to PK-*Insr*^w/w^ mice. **B**, Change in phospho-peptide abundance (Adj. p-values indicated in red) compared to the change in corresponding protein’s abundance between PK-*Insr*^f/f^ mice and PK-*Insr*^w/w^ mice. **C**, TOP: Schematic of phosphorylation sites on Cckar that were differentially abundant in any comparison. BOTTOM: The relative phospho-peptide abundance (arbitrary units) for each detected phospho-peptide for all samples and genotypes. **D,** Intracellular locations (outline created in Biorender) and functions of differentially abundant phospho-sites and their corresponding protein. The color in each attached phospho-site reflects the fold change, while the thickness of the border represents the Adj. p-value. Values are shown as mean ± SEM. *p<0.05, **p<0.01, ***p<0.001, ****p<0.0001 by background-based t-tests with p-values adjusted using Benjamini-Hochberg correction (C). See also Tables S3 and S5.

### Reducing insulin receptor signaling in acinar cells reduces fibrosis and inflammation

Our histological analyses, as well as our total proteomics and phosphoproteomics suggested that loss of the *Insr* in acinar cells prevented the formation of PanIN and prevented the induction of genes associated with pancreatic injury, such as Reg3a, Reg3b, and Reg2. To examine whether loss of *Insr* in acinar cells reduced the injury, inflammation, and/or fibrosis typically associated with Kras^G12D^ activation and PanIN formation, we imaged Sirius Red, alpha smooth muscle actin (αSMA), and F4/80 in 12-week-old pancreata (Figures 7A-B, S6A-D) to quantify the amount of collagen, activated fibroblasts, and macrophages present in the stroma. In PK-*Insr*^w/w^ control pancreata, Sirius Red, αSMA, and F4/80 staining surrounded the metaplastic ducts, containing ADM and PanIN, as well as the surrounding acini in nearby lobules (Figures 7A-B, S6A-D). In contrast, in the PK-*Insr*^f/f^ pancreata, the metaplasia that did form tended to be associated with less Sirius Red, αSMA, and F4/80 staining, as well as lower levels of staining in the neighboring lobules (Figures 7A-C, S6A-F). This supported our hypothesis that inflammation was reduced in the absence of *Insr* in Kras^G12D^-expressing acinar cells.

**Fig. 7.**
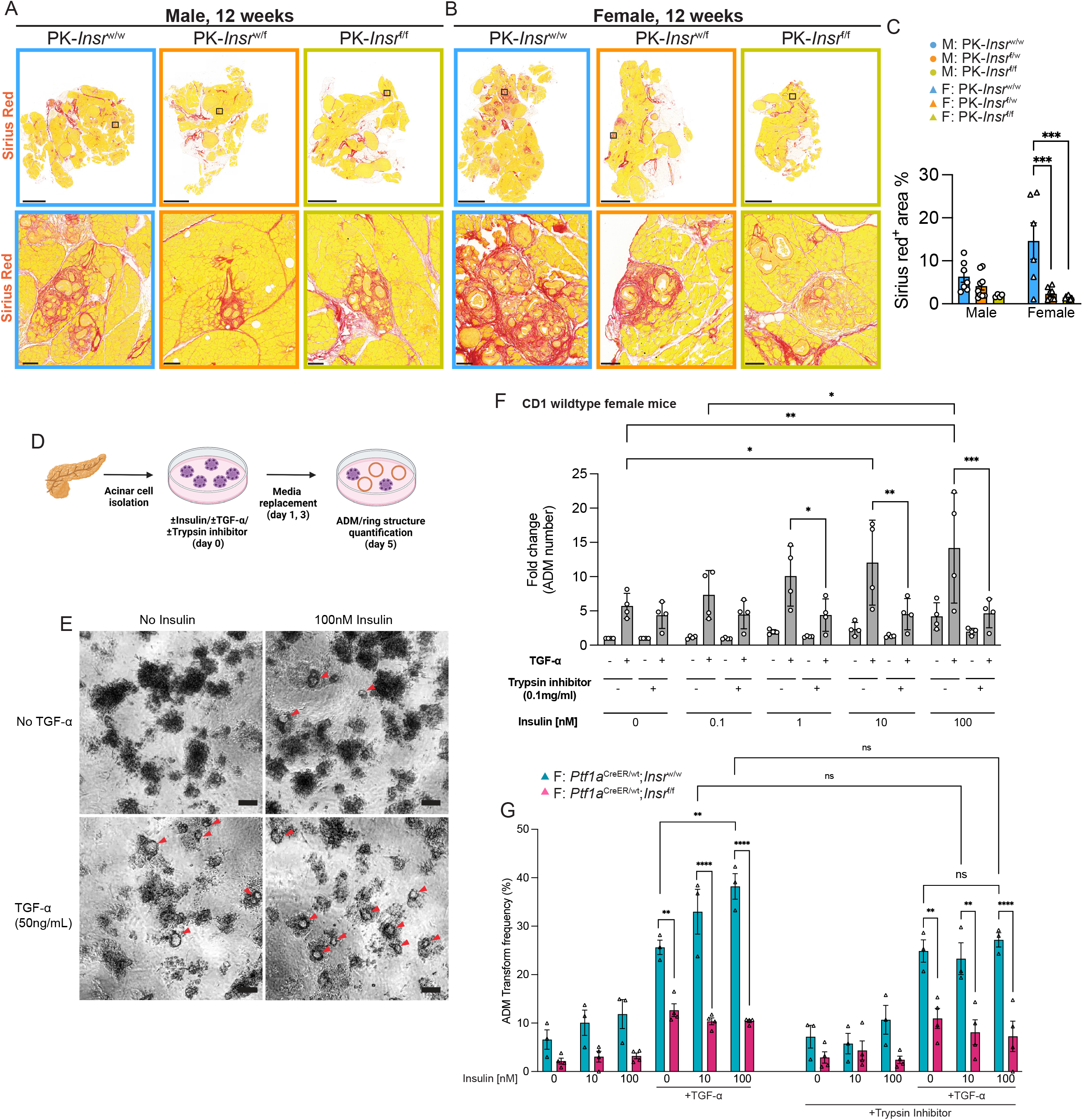
Insulin receptor signaling promotes PanIN initiation via increased inflammation associated with hyperactive digestive enzyme production. **A-B**, Representative whole section (top) and high-magnification (bottom) images of pancreatic slides stained with Sirius red from 12-week-old male (**A**) and female (**B**) PK-*Insr*^w/w^, PK-*Insr*^w/f^, and PK-*Insr*^f/f^ mice. **C,** Quantification of Sirius red positive area for mice from each genotype and sex (male (M:) or female (F:)) (n= 4-9). **D,** Schematic of the experimental design of the ADM formation assay using 3D acinar cell explant system. **E,** Representative bright field images of acinar cell clusters and duct formation (ring structures) on day 5 of treatment with or without 50 ng/mL TGF-α or 100 nM insulin in the absence of trypsin inhibitor. ADM structures are indicated by red arrows. **F,** Quantification of the fold change in ADM events in primary mouse acinar cell 3D explants (CD1 female) after 5 days of treatment with a combination of ± TGF-α and ± insulin and ± soybean trypsin inhibitor. (n = 4 independent experiments). Fold change values were calculated as treatment/negative control (no insulin or TGF-α or trypsin inhibitor) for each experiment. **G,** Quantification of ADM transformation rate of all acini clusters per well. Acinar clusters were isolated from female *Ptf1a*^CreER^;*Insr*^w/w^;nTnG (n=3) or *Ptf1a*^CreER^;*Insr*^f/f^;nTnG (n=4) mice. Acinar cell 3D explants were quantified after 5 days treatment with a combination of ± TGF-α and ± insulin with and without soybean trypsin inhibitor. Scale bars: 2 mm (**A**, **B**; top), 0.1 mm (**A**, **B**; bottom), and 100µm (**E**). Values are shown as mean ± SEM. *p<0.05, **p<0.01, ***p<0.001, ****p<0.0001 by one-way ANOVA (C), by 2-way ANOVA (F-G). See also Figures S6 and S7.

### Trypsin inhibitors block the effects of insulin on acinar-to-ductal metaplasia

Our proteomics data suggested reduced abundance of digestive enzymes in the absence of *Insr* in acinar cells in the context of Kras^G12D^ (Figure 5). Given that autoactivation of trypsinogen in acinar cells or in the pancreatic parenchyma may contribute to the induction of pancreatitis^41^, we reasoned that the reduced presence of enzymes in the absence of *Insr* could lessen tissue damage that spreads to other lobules. This would result in decreased ADM and PanIN formation, as in the PK-*Insr*^f/f^ pancreata. To test this hypothesis, we utilized an 3D *ex vivo* model of ADM formation from acinar cells (Figure 7D)^42,43^ to determine whether insulin levels affected the acinar to ductal transition. Acinar cells were isolated from wild-type male or female CD1 or Bl6 mice fed a normal chow diet. Wild-type female and male acinar cell clusters grown in collagen for 5 days predominantly maintained their acinus morphology, as previously shown (Figure 7E-CD1; male CD1 and Bl6 data not shown)^42^. TGF-α induced these clusters to form a duct-like lumen structure (Figure 7E-F-Cd1 female, Figure S7A-Bl6 female)^44^. Male acini typically did not form ring structures in response to TGF-α (Figure S7A-Bl6; CD1-data not shown), therefore we used female acini to test our hypothesis. Increasing concentrations of insulin significantly potentiated TGF-α induced ring formation (Figure 7E-F). Remarkably, the synergism between insulin and TGF-α was completely blocked by trypsin inhibitor (Figure 7F). Acini from *Ptf1a*^CreER^;*Insr*^f/f^;nTnG mice formed fewer ADM rings with or without TGF-α, and were unaffected by insulin (Figure 7G), demonstrating that these effects of insulin were mediated via *Insr*. Together, our data demonstrated that hyperinsulinemia-mediated trypsinogen pro-enzyme overproduction of, and subsequent autoactivation^45^, promotes local inflammation and initiation of PanIN lesions.

## DISCUSSION

The purpose of this study was to test the hypothesis that hyperinsulinemia in the obese state acts directly on acinar cells to promote pancreatic cancer initiation. Our data clearly support a model where Insr in acinar cells plays a causal role in supporting cancer initiation in the context of diet-induced obesity and mutant Kras. The contribution of direct insulin action on acinar cells during initiation from normal cells explains a large portion of the effects of obesity, but our results do not formally preclude minor roles for Insr in other local or distant cell types^10^. Our unbiased proteomic and phospho-proteomic analyses led us to propose a model in which diet-induced hyperinsulinemia acts directly on Insr in acinar cells to promote the physiological function of acinar cells, which is to supply digestive enzymes to break down lipid rich food in the duodenum. However, the increased production of enzymes increases the risk that more autoactivated trypsin will be present, as well. This increased propensity for trypsin-induced injury would result in sub-clinical levels of inflammation and acinar-to-ductal metaplasia. In the context of *Kras* mutations, this increased production can lead to increased chances of inflammation, which are known to enhance Kras^G12D^ signaling and promote an irreversible transformation in Kras^G12D^-expressing acinar cells^46^. Thus, our studies provide a key missing link explaining the connection between obesity and hyperinsulinemia and increased pancreatic inflammation and PanIN initiation.

Diet-induced obesity induces insulin hypersecretion, increases beta cell mass, and impairs insulin clearance resulting in sustained hyperinsulinemia^4,11^. Mice used in this study, which had the full complement of all 4 insulin alleles (*Ins1* and *Ins2*), exhibited higher fasting insulin levels (males 1000-1500 pmol/L; females 500 pmol/L) than our previous models with insulin gene dosage reduced to 1 or 2 copies (males, 400-800 pmol/L; females 100-200 pmol/L)^9,10^. The gradation of insulin levels between mice replete with insulin and those with reduced insulin production, is reflected in the severity of normal pancreas replacement by ductal metaplasia, PanIN, and tumors (e.g. this study PK-*Ins1*^+/+^:*Ins2*^+/+^ ∼90% replaced, PK-*Ins1*^+/+^;*Ins2*^-/-^ ∼25% replaced, PK-*Ins1*^-/-^;*Ins2*^+/+^ ∼1-4% replaced^9,10^). These findings reinforce that simply reducing insulin production limits PanIN initiation in the context of obesity.

All cells in the body have insulin receptors and require insulin signaling for key functions, including nutrient uptake for storage and anabolism. While the roles of Insr in hepatocytes, myocytes, and adipocytes are well studied^47–49^, the consequences of *Insr* loss in pancreatic acinar cells remain understudied. In this study, we specifically deleted the *Insr* gene from acinar cells using the *Ptf1a*^CreER^ allele and showed that insulin insensitivity in acinar cells had no obvious effects on systemic glucose homeostasis or serum insulin levels regardless of the *Kras* gene status. This suggests that the systemic regulation of glucose and insulin homeostasis are similarly perturbed in our mice fed HFD. Our data, combined with our previous studies^9,10^, effectively rule out an essential role for hyperglycemia in PanIN and PDAC formation, but do not preclude roles for hyperglycemia in the later stages of disease^50,51^.

Insr signaling activates mitogenic PI3K/AKT/mTOR and MAPK/ERK signaling cascades that are frequently mutated during tumorigenesis^52–54^, including in pancreatic cancer. Indeed, activating mutations in Kras, a key mediator of insulin and insulin-like growth factor signaling, drive the vast majority of pancreatic cancers^15^. Previous *in vitro* evidence supported the concept that hyperinsulinemia could promote cancer cell growth through over-activating the signaling cascades downstream of Insr protein^3,4,55,56^. Our findings show that insulin has a direct causal role in promoting the metaplastic changes needed for the acinar cell of origin to initiate tumorigenesis.

Mechanistically, our proteomic data demonstrate that *Insr* loss in acinar cells results in the coordinated reduction in digestive enzymes, with or without mutant Kras. Our observations confirm that insulin signaling normally supports exocrine function and are consistent with previous studies showing that amylase production is diminished by β-cell ablation using streptozotocin and restored with insulin injection^57–59^. The tendency towards reduced body weight in mice lacking acinar-cell Insr at some ages is also consistent with sub-clinical pancreatic insufficiency and a relative reduction in the ability to utilize ingested nutrients. Mutations that result in increased trypsin activity in the pancreas^8^, as well as animal studies using the Cck analog, caerulein, to stimulate enzyme secretion at supraphysiological levels^60^, support the concept that tight control of digestive enzyme function reduces the risk of PDAC. Our phospho-proteomic data identified hyper-phosphorylation of the Ccka receptor in *Insr*-knockout pancreas, linking local insulin signaling to Cck, a key endogenous regulator of acinar cell function and pancreatic weight^61–63^. It has been reported that HFD-associated inflammation can also promote islet Cck expression, which was proposed to play a role in obesity-associated PanIN formation^64^. Ultimately, our data indicate that hyperinsulinemia, acting through Insr, is the upstream driver of diet-induced inflammation via hyperactive digestive enzyme production. *Insr* loss in acinar cells counters the increased signal for acinar cell enzyme production or cell proliferation induced by HFD. Further studies are needed to fully understand the impact of dietary content on acinar cells and their susceptibility to Kras^G12D^-mediated transformation.

Our proteomic analyses revealed other key mechanisms associated with suppressed pancreatic cancer initiation from acinar cells lacking *Insr*. For example, mRNA splicing and translation factors, known targets of insulin signaling via mTor^48^, are differentially abundant in the pancreas after *Insr* loss in Kras^G12D^-expressing acinar cells, suggesting mechanisms by which insulin may regulate digestive enzyme production. We also found evidence that reduced insulin signaling affects cellular metabolism. Previous studies suggested that insulin promotes glycolysis in wildtype acinar cells to protect them during pancreatitis^65^, however, detailed analyses of cell-type-specific mechanisms await single-cell proteomic and single-cell metabolomic characterization of this model.

In summary, our data strongly suggest that insulin receptor signaling in acinar cells contributes to the PanIN and PDAC development. Our data illustrate the complex and interconnected molecular mechanisms by which hyperinsulinemia, acting directly through acinar cell Insr, promotes early pancreatic tumourigenesis. Thus, we can infer that targeting insulin receptor signaling pathways, or hyperinsulinemia itself, may be beneficial in treating and preventing pancreatic cancer in patients. The ligand of this pathways, insulin, is one of the most clinically actionable hormones in our bodies and it can be easily measured and can be modulated by diet, exercise, as well as FDA-approved drugs. Thus, our novel observations that modifying insulin alone can reduce the risk of initiating PDAC elucidate multiple prevention strategies that could be trialed in humans.

### Limitations of the Study

While our study breaks new ground, we also acknowledge its limitations and scope. Our results led to a focus on cancer initiation, so we cannot extrapolate these findings and models to our understanding of advanced tumor progression or metastasis. Futures studies will be needed to dissect the role of hyperinsulinemia and insulin receptors on these later stages.

Some of our data, including non-significant trends, are suggestive of sex differences in the role of Insr mechanisms and/or interactions between PDAC initiation and whole-body glucose homeostasis. Additional work, beyond the scope of this study, will be required to investigate this. We think it is unlikely that the mechanistic role of Insr is fundamentally different between sexes, although it seems possible that inherent differences in male and female acini affect their predisposition to undergo metaplasia in the mouse.

A technical limitation of our study, and the field, is that existing antibody reagents are not specific enough to perform accurate anti-Insr staining that would allow us to determine whether acinar cells that contributed to PanIN formation did so due to incomplete recombination at the *Insr* floxed allele. Moreover, coverage of the proteome was not complete and biased against membrane proteins. For example, analysis of the abundance and phosphorylation of Igf1r, a protein that may compensate in *Insr* knockout cells, will require targeted assays and/or membrane fractionation. Insulin could bind to homodimeric Igf1 receptors and heterodimeric Insr/Igf1r hybrids, although at a lower affinity^21^, to mediate its pro-tumourigenic effects. Future studies on acinar cell-specific *Igf1r* knockout and/or *Insr*/*Igf1r* double knockout would be necessary to delineate any *Igf1r* contribution to PanIN formation. Nevertheless, it is clear that simply reducing *Insr* gene dosage was sufficient to reduce the effects of hyperinsulinemia on PanIN initiation.

## METHODS

### Mouse models

All animal experiments were conducted at the University of British Columbia with approval of the University of British Columbia Animal Care Committee in accordance with Canadian Council for Animal Care guidelines. All alleles have been described previously^24,66–70^. *Kras*^LSL-G12D/w^ (#008179), *Insr*^f/f^ (#006955), and nuclear TdTomato-to-nuclear EGFP (nTnG) mice (#023035) were purchased from Jackson Labs (Bar Harbour, USA). *Ptf1a^CreER/w^* mice (C57BL/6) were a gift from Chris Wright (Vanderbilt, USA). Mice were maintained on C57BL/6 genetic background and housed at the University of British Columbia Modified Barrier Facility which was temperature-controlled (22°C) and specific pathogen-free. They were kept on a 12:12 hr light: dark cycle with food and drinking water *ad libitum*. To generate genetic background-matched *Ptf1a^CreER/w^;Kras^LSL-G12D/w^;Insr^w/w^;nTnG*, *Ptf1a^CreER/w^;Kras^LSL-G12D/w^;Insr^w/f^;nTnG*, and *Ptf1a^CreER/w^;Kras^LSL-G12D/w^;Insr^f/f^;nTnG* mice, *Ptf1a^CreER/w^;Kras^LSL-G12D/w^;Insr^w/f^;nTnG* mice were bred with *Insr^w/f^* mice. After weaning (3 weeks), the resulting litters were fed with high fat diet (HFD), and at 4 weeks of age, recombination was induced over three consecutive days by subcutaneous injections of tamoxifen in corn oil (20 mg/mL) of 5 mg tamoxifen/40 g body mass. Both male and female mice were used in this study and humane endpoint was met in this study by losing 20% of the max body weight. One cohort of mice was euthanized at 10 months of age for histopathology analyses and the whole pancreas was used for histopathology analyses. Another cohort of mice was euthanized at 12 weeks of age. For mice euthanized at 12 weeks, each pancreas was cut into three pieces roughly based on the pancreas anatomical structure: head, body, and tail. The body piece was used for histopathological analyses, while the other pieces were snap frozen in liquid nitrogen and kept at –80 °C. The head piece was processed for proteomics and phospho-proteomics analyses and part of the tail was used for RNA analyses (see below).

### Assessment of glucose homeostasis

Mouse body weight and fasting blood glucose levels were measured every 4 weeks and fasting insulin were measured every 3 months. Before the measurements, mice were fasted for 4 hours in clean and fresh cages during the light period. One drop of blood was collected from each mouse’s saphenous vein and a Lifescan OneTouch Ultra Mini glucometer was used to measure the fasting blood glucose levels. About 30 µl of blood was collected with a heparinized microhematocrit capillary tube (Fisher Scientific, 22-362566, Waltham, MA, USA) for measuring fasting insulin levels. The collected blood was centrifuged at 10,000 rpm for 10 minutes to collect the blood serum. Then the blood serum was kept at

-20°C until used to measure the fasting insulin levels with insulin ELISA (80-INSMSU-E10; ALPCO Diagnostics, Salem, NH).

### Histopathological, morphological, and immunohistochemical analyses

Pancreata were fixed in 4% paraformaldehyde for 24 hours at 4°C followed by paraffin embedding. Mouse pancreata were sectioned then stained with hematoxylin and eosin (H&E), anti-GFP immunohistochemistry, or Alcian blue, as described^9,10,71^. The stained slides were scanned with a 20× objective using a 3DHISTECH Panoramic MIDI (Quorum Technologies Inc. Guelph, Canada) slide scanner. One 12-week-old mouse with a GFP labeling efficiency of acinar cells below 20% was excluded from further analyses. Histopathological analyses were conducted in a de-identified manner and verified separately by J.L.K and D.F.S. All histopathological analyses were performed on one of the stained sections that displayed the maximal pancreatic cross-sectional area unless otherwise stated. Every gland with a lumen was categorized as normal, ADM, PanIN, or neoplasia, and glands representing more than one of these categories were scored based on their highest-grade feature. The total pancreatic area, PanIN plus tumor area, PanIN area, or normal acinar cell area were measured as previously described^9,10^. Briefly, the total pancreatic area, PanIN plus tumor area, PanIN area, or normal acinar cell area was determined by masking all pancreatic tissue, selective masking of the PanIN plus tumor area, selective masking of the only PanIN area, or selective masking of the normal acinar cell area by Adobe Photoshop. Pixels for the total pancreatic area or each histological feature were measured by ImageJ and this was used to calculate the percentage area occupied by each histological feature. For Alcian blue positive area, Adobe Photoshop 2020 Black & White function was used to highlight the blue area (red filter). The total pixels for pancreas or Alcian blue positive area were counted using ImageJ. For Sirius Red positive area, Adobe Photoshop 2020 Black & White function was used to highlight the red area (green filter). The total pixels for pancreas or Sirius Red positive area were counted using ImageJ. Immunohistochemistry and immunofluorescence staining was performed according to published standard protocols^10,24,71^. Primary antibodies were rabbit anti-cytokeratin 19 (Ck19) (Abcam, ab133496, 1:1000), mouse αSMA (Sigma Aldrich, A2547, 1:100) and goat anti-GFP (Abcam, ab6673, 1:200). Secondary antibodies were Cy3 conjugated AffiniPure Donkey Anti-Rabbit IgG (Jackson ImmunoResearch, 711-165-152, 1:500), biotin-conjugated donkey anti-rabbit (Jackson ImmunoResearch, 711-065-152, 1:500), or biotin-conjugated rat anti-goat (Vector Laboratories, MP-7404, 1:2). Slides were scanned with a 20× objective using a 3DHISTECH Panoramic MIDI slide scanner. Ck19, Cpa1, pErk, pAkt, or F4/80 positive areas or αSMA positive area was measured with Adobe Photoshop 2020 using the Black & White function to filter and highlight the brown area (blue filter) or the red fluorescence area, respectively, which was then taken as a percent of pixels in the pancreatic section.

### RNA isolation and analysis

A piece of the tail from pancreata collected from female mice at 12 weeks of age (PK-*Insr*^w/w^ (n=4), PK-*Insr*^w/f^ (n=3), PK-*Insr*^f/f^ (n=4)), were frozen with liquid nitrogen and shattered with a C01 Manual Pulverizer and RNA was isolated with RNeasy Micro Kit (QIAGEN, 74004). After RNA extraction, the Agilent 2100 Bioanalyzer was used to perform sample quality control. Samples that qualified (RIN>5) were prepped using the standard protocol for Illumina Stranded mRNA prep (Illumina). Sequencing was done using the Illumina NextSeq 2000 with paired-end 59 bp × 59 bp reads. Sequencing data were demultiplexed using Illumina’s BCL Convert and aligned to the Mus Musculus (mm10) reference sequence using DRAGEN RNA app on Basespace Sequence Hub (https://support-docs.illumina.com/SW/DRAGEN_v41/Content/SW/DRAGEN/TPipelineIntro_fDG.htm).

Differential expression analysis was done using DESeq2 R package^72^. A low-count filter was applied to remove genes with < 5 raw reads in > 75% of samples. Genes were normalized by variance stabilizing transformation and deemed to be differentially expressed when their Benjamini & Hochberg adjusted p-values were < 0.05. Gene Set Enrichment Analysis (GSEA) was done by the clusterProfiler R package^73^, using the MSigDB Hallmark gene set knowledgebase^74,75^. The sign of log_2_ fold change * (–log_10_ p-value) was used as the rank score in GSEA. The “tags” values of the GSEA output were plotted as “Gene Ratio”, which is the ratio of the number of leading-edge genes (genes that drove the enrichment) to the total number of genes in each gene set. The results of differential expression or GSEA were plotted in a volcano plot or dot plot using ggplot2 R package^75^. The code for RNA-seq analyses is available at https://github.com/hcen/PDAC_InsrKO. The sequencing data can be accessed through the GEO repository: GSE229053.

### Proteomics analyses

The whole head of pancreata collected from female mice at 12 weeks of age (*Ptf1a*^CreER^;*Kras*^LSL-G12D^;*Insr*^w/w^;nTnG (n=3), *Ptf1a*^CreER^;*Kras*^LSL-G12D^;*Insr*^w/f^;nTnG (n=6), *Ptf1a*^CreER^;*Kras*^LSL-G12D^;*Insr*^f/f^;nTnG (n=6), *Ptf1a*^CreER^;*Insr*^w/w^;nTnG (n=3), and *Ptf1a*^CreER^;*Insr*^f/f^;nTnG (n=2) mice) were frozen and used for (phospho)proteomic analyses. The frozen sample was ground into powder in a liquid nitrogen-cooled mortar and pestle and kept on dry ice until mass-spec analysis. Proteins were extracted from cryopulverized tumors in a buffer containing 5% sodium dodecyl sulfate (SDS) and 100 mM TRIS pH 7.8 supplemented with PhosStop phosphatase inhibitor cocktail (Millipore-Sigma, part# 4906845001). The protein concentration of the lysate was determined using bicinchoninic acid assay (BCA) (Thermo Fisher Scientific, Waltham, MA, USA, part# 23225), and protein disulfide bonds were reduced and free Cysteines alkylated with 20 mM tris (2-carboxyethyl)phosphine (TCEP) and 30 mM iodoacetamide, respectively (Millipore-Sigma, part# 646547, and l1149 respectively). Proteolytic digestion of 250 µg of total protein was performed using S-TRAP Mini columns (Protifi LLC, Huntington NY, USA)^76^. Resultant tryptic peptides were then vacuum concentrated and desalted using Oasis HLB SPE cartridges (Waters, Milford, MA, USA, part# WAT094225). Peptides were vacuum concentrated and reconstituted in 0.1% trifluoro acetic acid (TFA), and 10% of the total sample was reserved for measurement of the total proteome. The remaining 90% of the sample was diluted in 80% acetonitrile with 0.1% TFA for automated phosphopeptide enrichment using AssayMap Fe-NTA (III) immobilized metal affinity chromatography (IMAC) cartridges and a Bravo liquid handling system using the phosphopeptide enrichment v 2.1 application (Agilent Technologies, Santa Clara, CA, USA, part# G5496-60085).

### LC-MS/MS acquisition and data analysis

Samples for both total proteomics and phosphoproteomics were analyzed by data dependent acquisition (DDA) using an Easy-nLC 1200 and Q Exactive Plus (both Thermo Fisher Scientific, Waltham, MA, USA). Samples were first loaded onto a precolumn (Acclaim PepMap 100 C18, 3 µm particle size, 75 µm inner diameter x 2 cm) in 0.1% formic acid (buffer A), and gradient elution was performed using a 100-min method from 3 to 40% buffer B (84% acetonitrile, 0.1% formic acid) on the analytical column (Acclaim PepMap 100 C18, 2 µm particle size, 75 µm inner diameter x 25 cm) at a flow rate of 300 nL/min. MS scans were acquired between 350-1,500 m/z at a resolution of 70,000, with an automatic gain control (AGC) target of 1 x 10e610E6 ions and a maximum injection time of 50 ms. The top 15 precursor ions with charge states +2, +, +3, and +4 were isolated with a window of 1.2 m/z, an AGC target of 2 x 10e410E4 and a maximum injection time of 64 ms and fragmented using a normalized collision energy (NCE) of 28. MS/MS were acquired at a resolution of 17,500 and the dynamic exclusion was set to 30 s. DDA MS raw data was processed with Proteome Discoverer 2.5 (Thermo Fisher Scientific, Waltham, MA, USA) and searched using Sequest HT against the mouse reference proteome FASTA database from Uniprot (downloaded October 1^st^ 2021; 17,054 forward sequences). The enzyme specificity was set to trypsin with a maximum of 2 missed cleavages. Carbamidomethylation of cysteine was set as a fixed modification and oxidation of methionine, as well as phosphorylation of serine, threonine, and tyrosine as variable modifications. The precursor ion mass tolerance was set to 10 parts per million, and the product ion mass tolerance was set to 0.02 Da. Percolator was used to assess posterior error probabilities and the data was filtered using a false discovery rate (FDR) of <1% on peptide and protein level. The Minora node of Proteome Discoverer was used for label free quantitation. For the human comparison, Pathway enrichment was performed on our mouse data and PDAC human data^34^ using Reactome (v80)^77^. We performed differential abundance analysis on the human data using the R package limma^78^. For our data, we used a cutoff of Adj. p<0.05. For the human data, we used Adj. p<1^-19^ and 1^-12^ for the total proteins and phospho-proteins respectively to ensure a comparable proportion of the total number detected. The mass spectrometry proteomics data have been deposited to the ProteomeXchange Consortium via the PRIDE^79^ partner repository with the dataset identifier PXD033845. All code is available at https://github.com/johnsonlabubc/PDAC-Insr-KO-Total-and-Phosphoproteomic-Analysis.

### Primary mouse acinar cell isolation and three-dimensional culture

Primary pancreatic acinar cells were isolated from pancreata of wild-type female or male mice of the indicated mouse background fed normal chow diet (LabDiet Rodent Diet 20 Cat#5053) at the age of 6-8 weeks. *Ptf1a*^CreER^*;Insr*^f/f^;nTnG mice and their control mice (*Ptf1a*^CreER^;*Insr*^w/w^;nTnG) were given tamoxifen via subcutaneous injection two weeks prior to cell isolation. The acinar cell isolation procedure was adapted from the protocol described by Martinez and Storz^43^. Briefly, the mouse pancreas was harvested and washed three times in 1× HBSS and then minced into 1-5 mm pieces. The fragmented tissue was then digested with 5 mL (0.4 mg/mL) collagenase P solution (Sigma, Cat. No. 11213857001) in RPMI1640 (Sigma, Cat. No. R0883) at 37°C with gentle shaking for 18 minutes. HBSS (10 mL) with 5% FBS was added to terminate the digestion reaction, followed by 3 washes with 10 mL HBSS (5% FBS) to remove the residual collagenase P. After each wash, the tissue was pelleted (450g RCF, 2 min at room temperature) and the supernatant was removed. The digested tissue was resuspended in 10 mL HBSS (5% FBS) and filtered through a 100 μm cell strainer, followed by a wash with 10 mL HBSS (5% FBS). The filtrate was gently added to 20 mL HBSS 30% FBS cushion to form layers of cells. The cell mixture was centrifuged at 180g RCF at room temperature for 2 minutes to pellet acinar cells. Appropriate number of acinar cells were then resuspended in premade collagen solution (1 mg/ml rat tail type1 collagen (VWR, Cat. No. CACB354236), 10× Waymouth’s media (Cedarlane Labs, Cat. No. W1001-03), RPMI1640 complete media (1% FBS, 1× penicillin/streptomycin (VWR, Cat. No. CA45000-650), 1 μg/mL dexamethasone (Sigma, Cat. No. D2915), adjusted to pH = ∼7.8 with 1 M NaOH) and plated (50 μL per well) in collagen pre-coated 96-well plates. After solidification, 100 μL of the RPMI1640 complete media with or without insulin (0.1 nM-100 nM) (Sigma, Cat. No. 11061-68-0), TGF-α (Fisher Scientific, Cat. No. PHG005) (50ng/mL) or combined treatments, with or without 0.1 mg/mL soybean trypsin inhibitor (Sigma, Cat. No. T9128) was added. *Ptf1a*^CreER^*;Insr*^f/f^;nTnG acinar cells and the control cells were cultured with Human plasma-like media (Gibco, Cat. No. A4899102), instead of RPMI1640, to ensure the survival of *Insr-*deficient acinar cells. Media was replaced on day 1 and 3, and the numbers of ADM events in each well were quantified on day 5. Isolated cells were maintained at 37°C in a humidified incubator with 5% CO_2_.

## QUANTIFICATION AND STATISTICAL ANALYSIS

Animals were excluded from the histopathological analyses if they were found dead. Statistical analyses were conducted with GraphPad Prism 9.3.0. The Shapiro-Wilk test was used to assess data normality. One-way ANOVA was performed unless otherwise stated. Mixed-effect analyses were run for glucose homeostasis data (mouse fasting glucose level, and fasting insulin level) and a compare groups of growth curves permutation test for body weight gains^80^. When comparing the histopathological measurements between male and female mice from the same genotype, a two-tailed student’s t-test was run for normally distributed data and a Mann-Whitney test was performed for non-normally distributed data. Statistical parameters, including the exact value of sample size n (animal number), precision measures and dispersion (mean ± SEM), and statistical significance levels, are reported in the Figures and Figure legends. A p-value <0.05 was considered significant. In Figures, asterisks denote statistical significance (*p<0.05, **p<0.01, ***p<0.001, and ****p<0.0001).

For total proteomics data, normalization, imputation, and differential expression analyses were performed on the protein abundances using Proteome Discoverer. Differential expression analysis used background-based t-tests with p-values adjusted using Benjamini-Hochberg correction. The proteins were filtered for significant comparisons (Adj. p<0.05), and heatmaps were created with resultant protein lists. We also applied k-means clustering and extracted clusters with patterns of interest^81^. From this heatmap, proteins were identified and values for all samples were plotted in a heatmap and sorted with hierarchical clustering. We used STRING (v11) to generate an edge table with differentially expressed total proteins (Adj. p<0.05) with an absolute log_2_(Fold Change) cut-off of 2.5 in *Ptf1a*^CreER^;*Kras*^LSL*-*G12D^;*Insr*^w/w^;nTnG compared to *Ptf1a*^CreER^;*Kras*^LSL-G12D^;*Insr*^f/f^;nTnG. Protein-protein relationships were only included if they involved experimental evidence or interactions annotated in other databases^35^. Cytoscape (v3.9.1) was used to visualize the network^82^, with background images created with BioRender.com.

Phospho-proteomics data were analyzed two ways. In the first approach, we only used phospho-peptides with measurements made in a minimum of 2/3 of all the samples and at least one sample per group. We imputed missing phospho-protein values in samples before normalization and statistical analysis using the same process as the total proteomics, allowing a broader assessment of the phospho-protein landscape. These were plotted using volcano plots and against the corresponding proteins in the total protein data set with calculated log_2_(Fold Change) of the median. In the second approach, we aggregated the abundances of each sample for each phospho-site detected, and normalized each of the detected phospho-sites to the total abundance of the corresponding protein. We only included phospho-sites where every sample had an experimentally measured total protein value. The second method used differential expression analysis performed using the R package limma on the *Ptf1a*^CreER^;*Kras*^LSL-G12D^;*Insr*^w/w^;nTnG versus *Ptf1a*^CreER^;*Kras*^LSL-G12D^;*Insr*^f/f^;nTnG comparison^78^. Phosphosite^38^ was used to determine upstream activators, cellular location, and function of each protein, as well as the specific kinases of the detected phospho-sites, if known. STRING was used to identify proteins that interact with each other^35^.

## Acknowledgements

The authors thank members from Johnson and Kopp labs for discussions. We thank Christopher Wright (Vanderbilt, USA) for the *Ptf1a*^CreER^ mice. RNA sequencing analysis was performed with the help of the SBME-Seq facility at UBC. Bioinformatics support was provided by Stephane Flibotte at the UBC Life Sciences Bioinformatics Core.

