## Supplemental Figures for "Hyperinsulinemia acts via acinar insulin receptors to initiate pancreatic cancer by increasing digestive enzyme production and inflammation"

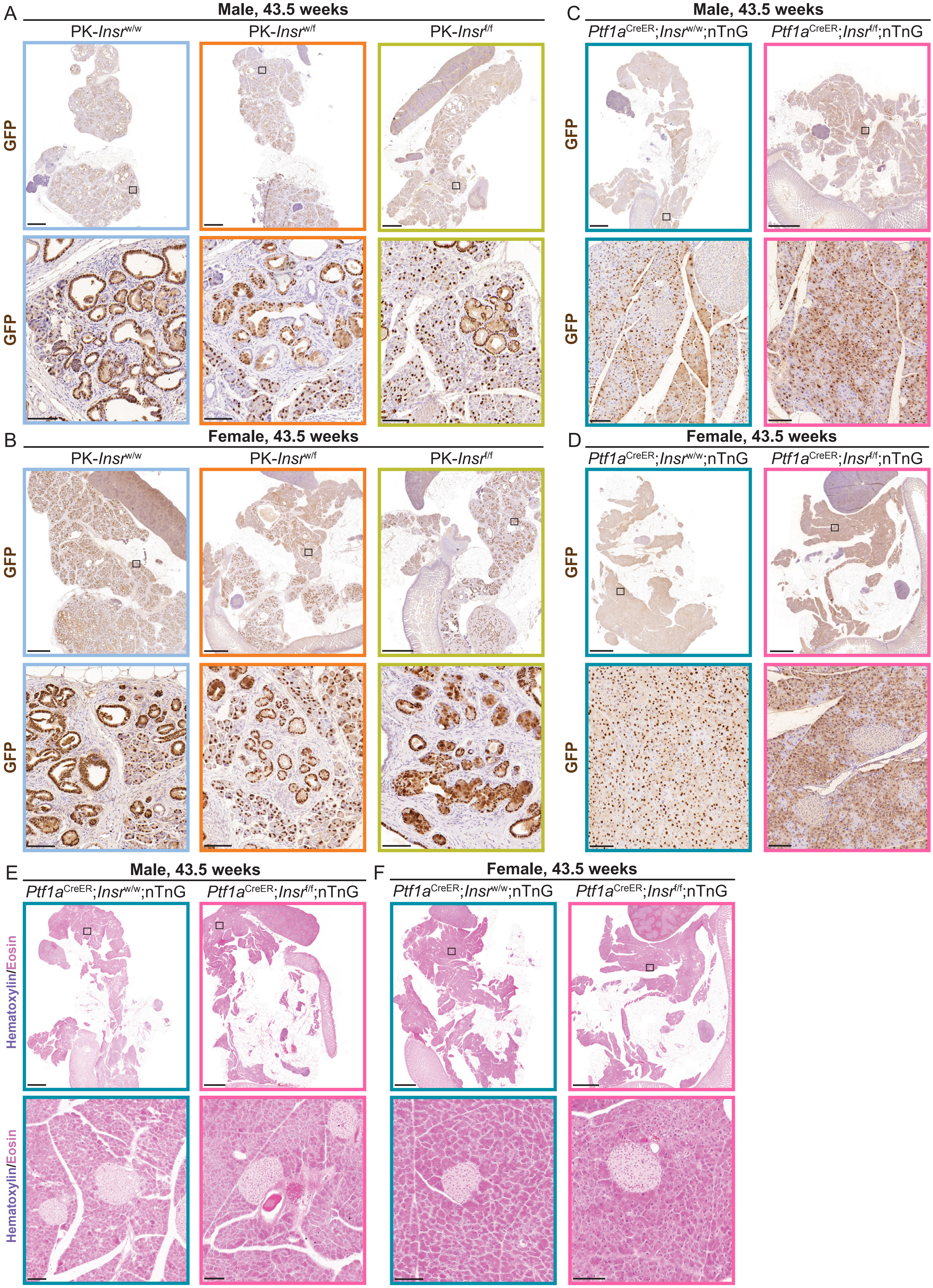
A
Male, 43.5 weeks
PK-Insrw/w
PK-Insrw/f
PK-Insrf/f
GFP
GFP
B
Female, 43.5 weeks
PK-Insrw/w
PK-Insrw/f
PK-Insrf/f
GFP
GFP
C
Male, 43.5 weeks
Ptf1aCreER;Insrw/w;nTnG
Ptf1aCreER;Insrf/f;nTnG
GFP
GFP
D
Female, 43.5 weeks
Ptf1aCreER;Insrw/w;nTnG
Ptf1aCreER;Insrf/f;nTnG
GFP
GFP
E
Male, 43.5 weeks
Ptf1aCreER;Insrw/w;nTnG
Ptf1aCreER;Insrf/f;nTnG
Hematoxylin/Eosin
Hematoxylin/Eosin
F
Female, 43.5 weeks
Ptf1aCreER;Insrw/w;nTnG
Ptf1aCreER;Insrf/f;nTnG

**Figure S1. The *Ptf1a*$^{CreER}$ allele labeled acinar cells and PanINs in 43.5-week-old mice, related to Figure 1.**

**A-B,** Representative whole section (top) and high-magnification (bottom) images of immunohistochemical staining for GFP expressed from nTnG lineage tracing allele in 43.5-week-old male (**A**) and female (**B**) PK-*Insr*$^{w/w}$, PK-*Insr*$^{w/f}$, and PK-*Insr*$^{f/f}$ mice. **C-D**, Representative whole section (top) and high-magnification (bottom) images for immunohistochemical staining for GFP expressed from nTnG lineage tracing allele in 43.5-week-old male (**C**) and female (**D**) *Ptf1a*$^{CreER}$;*Insr*$^{w/w}$;nTnG and *Ptf1a*$^{CreER}$;*Insr*$^{f/f}$;nTnG mice. **E-F**, Representative whole section (top) and high-magnification (bottom) H&E images of pancreatic slides from 43.5-week-old male (**E**) and female (**F**) *Ptf1a*$^{CreER}$;*Insr*$^{w/w}$;nTnG and *Ptf1a*$^{CreER}$;*Insr*$^{f/f}$;*nTnG* mice. Scale bars: 2 mm (top) and 0.1mm (bottom).

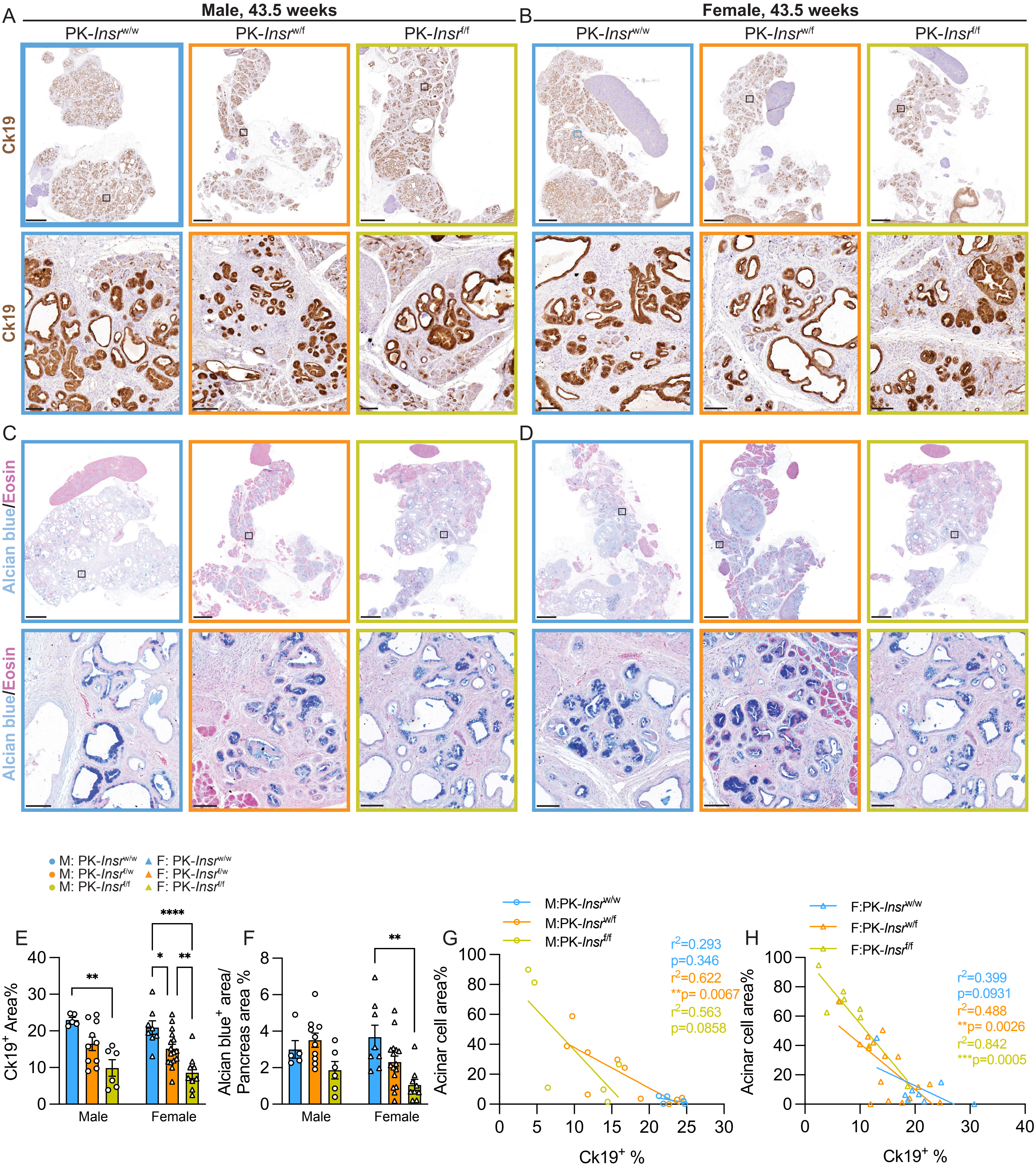

A
Male, 43.5 weeks
B
Female, 43.5 weeks
PK-Insr w/w
PK-Insr w/f
PK-Insr f/f
Ck19
C
D
Alcian blue/Eosin
M: PK-Insr w/w
M: PK-Insr f/w
M: PK-Insr f/f
F: PK-Insr w/w
F: PK-Insr f/w
F: PK-Insr f/f
E
Ck19+ Area%
Male
Female
F
Alcian blue+ area/ Pancreas area %
G
M:PK-Insr w/w
M:PK-Insr w/f
M:PK-Insr f/f
r2=0.293
p=0.346
r2=0.622
**p= 0.0067
r2=0.563
p=0.0858
Acinar cell area%
Ck19+ %
H
F:PK-Insr w/w
F:PK-Insr w/f
F:PK-Insr f/f
r2=0.399
p=0.0931
r2=0.488
**p= 0.0026
r2=0.842
***p=0.0005

**Figure S2. Loss of *Insr* in acinar cells reduced ductal metaplasia and Alcian blue$^+$ PanIN lesions, related to Figure 1.**

**A-B,** Representative whole section (top) and high-magnification (bottom) images of immunohistochemical staining of Ck19 for male (**A**) and female (**B**) PK-*Insr*$^{w/w}$, PK-*Insr*$^{w/f}$, and PK-*Insr*$^{f/f}$ mice. **C-D,** Representative whole section (top) and high-magnification (bottom) images of pancreatic slides from male (**C**) and female (**D**) PK-*Insr*$^{w/w}$, PK-*Insr*$^{w/f}$, and PK-*Insr*$^{f/f}$ mice stained with Alcian blue. **E,** Quantification of Ck19$^+$ area for mice from each genotype and sex (M or F) (n= 5-15). **F,** Quantification of Alcian blue$^+$ area for mice from each genotype and sex (M or F) (n= 5-16). **G-H,** The correlation of acinar cell area and Ck19$^+$ area for male (**G**) and female (**H**) mice. The maximum value for Ck19$^+$ area was ~20-30% due to stromal expansion in the parenchyma around the Ck19$^+$ area. Scale bars: 2 mm (top) and 0.1 mm (bottom). Values are shown as mean ± SEM. *$p<0.05$, **$p<0.01$, ***$p<0.001$, ****$p<0.0001$ by one-way ANOVA (E), Kruskal-Wallis test (F), or by correlation test (G-H).

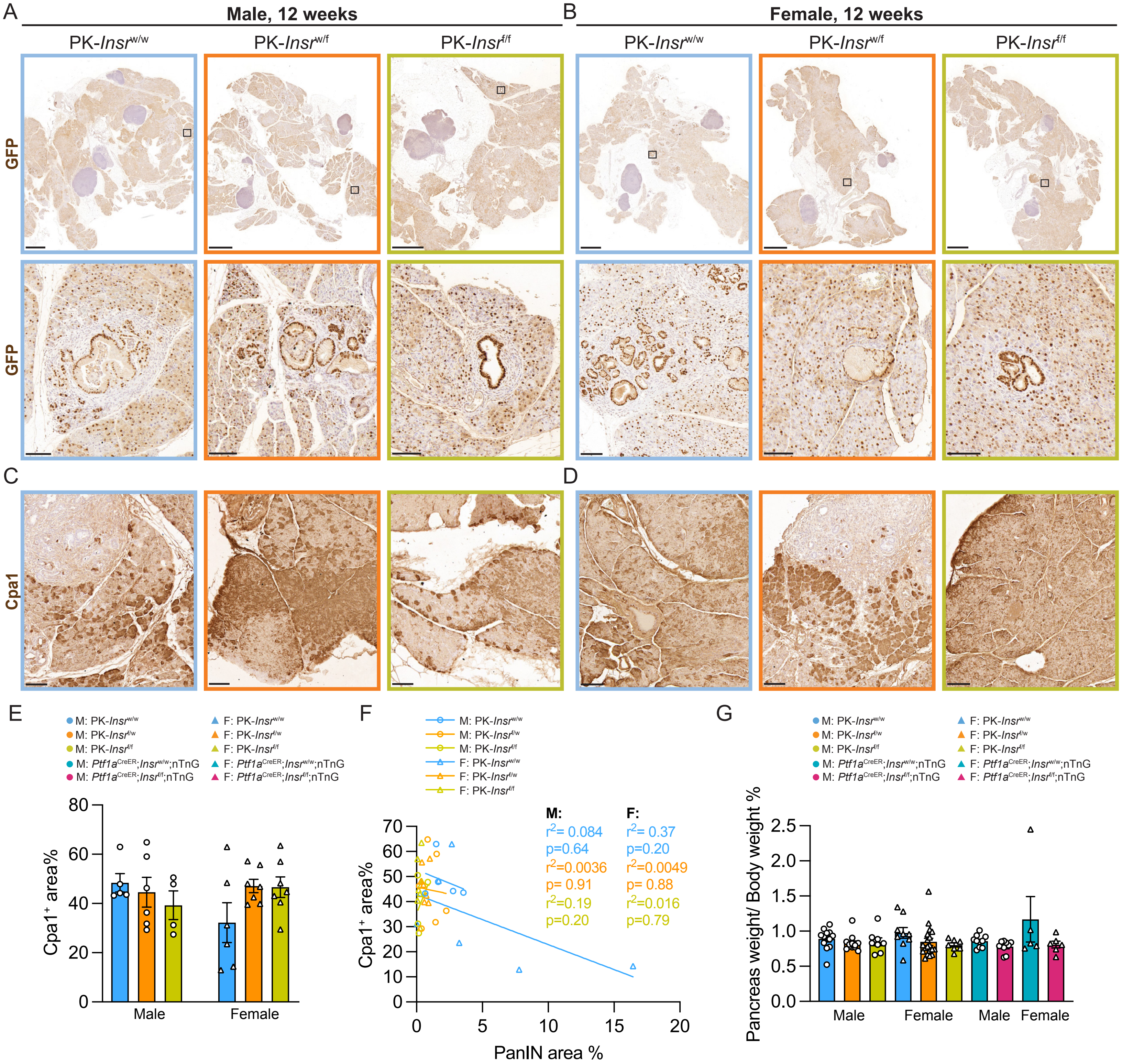

A
Male, 12 weeks
B
Female, 12 weeks
PK-Insr w/w
PK-Insr w/f
PK-Insr f/f
GFP
GFP
C
D
Cpa1
E
M: PK-Insr w/w
M: PK-Insr f/w
M: PK-Insr f/f
M: Ptf1a CreER;Insr w/w;nTnG
M: Ptf1a CreER;Insr f/f;nTnG
F: PK-Insr w/w
F: PK-Insr f/w
F: PK-Insr f/f
F: Ptf1a CreER;Insr w/w;nTnG
F: Ptf1a CreER;Insr f/f;nTnG
Cpa1+ area%
Male
Female
F
M: PK-Insr w/w
M: PK-Insr f/w
M: PK-Insr f/f
F: PK-Insr w/w
F: PK-Insr f/w
F: PK-Insr f/f
M:
r2= 0.084
p=0.64
r2=0.0036
p= 0.91
r2=0.19
p=0.20
F:
r2= 0.37
p=0.20
r2=0.0049
p= 0.88
r2=0.016
p=0.79
Cpa1+ area%
PanIN area %
G
Pancreas weight/ Body weight %
Male
Female
Male
Female

**Figure S3. *Insr* loss does not affect pancreatic weight or Cpa1$^{+}$ area in 12-week-old mice, related to Figure 3.**

A-B, Representative whole section (top) and (bottom) images of immunohistochemical staining for GFP expressed from the nTnG lineage tracing allele in 12-week-old male (**A**) and female (**B**) PK-*Insr*$^{w/w}$, PK-*Insr*$^{w/f}$, and PK-*Insr*$^{f/f}$ pancreata. **C-D,** Representative high-magnification images of immunohistochemical staining of Cpa1 for male (**C**) and female (**D**) PK-*Insr*$^{w/w}$, PK-*Insr*$^{w/f}$, and PK-*Insr*$^{f/f}$ mice. **E,** Quantification of Cpa1$^{+}$ area for mice from each genotype and sex (M or F) (n= 4-7). **F,** The correlation of PanIN area and Cpa1$^{+}$ area for each genotype and sex (M or F) mice. **G,** The ratio of pancreatic weight to mouse body weight for male (M) and female (F) *Ptf1a*$^{CreER}$;*Insr*$^{w/w}$;nTnG, *Ptf1a*$^{CreER}$;*Insr*$^{f/f}$;nTnG mice, PK-*Insr*$^{w/w}$, PK-*Insr*$^{w/f}$, and PK-*Insr*$^{f/f}$ mice (n=3-15). Scale bars: 2 mm (top) and 0.1 mm (bottom). Values are shown as mean ± SEM. Statistical tests were performed with one-way ANOVA (E, G), or by simple linear regression (F).

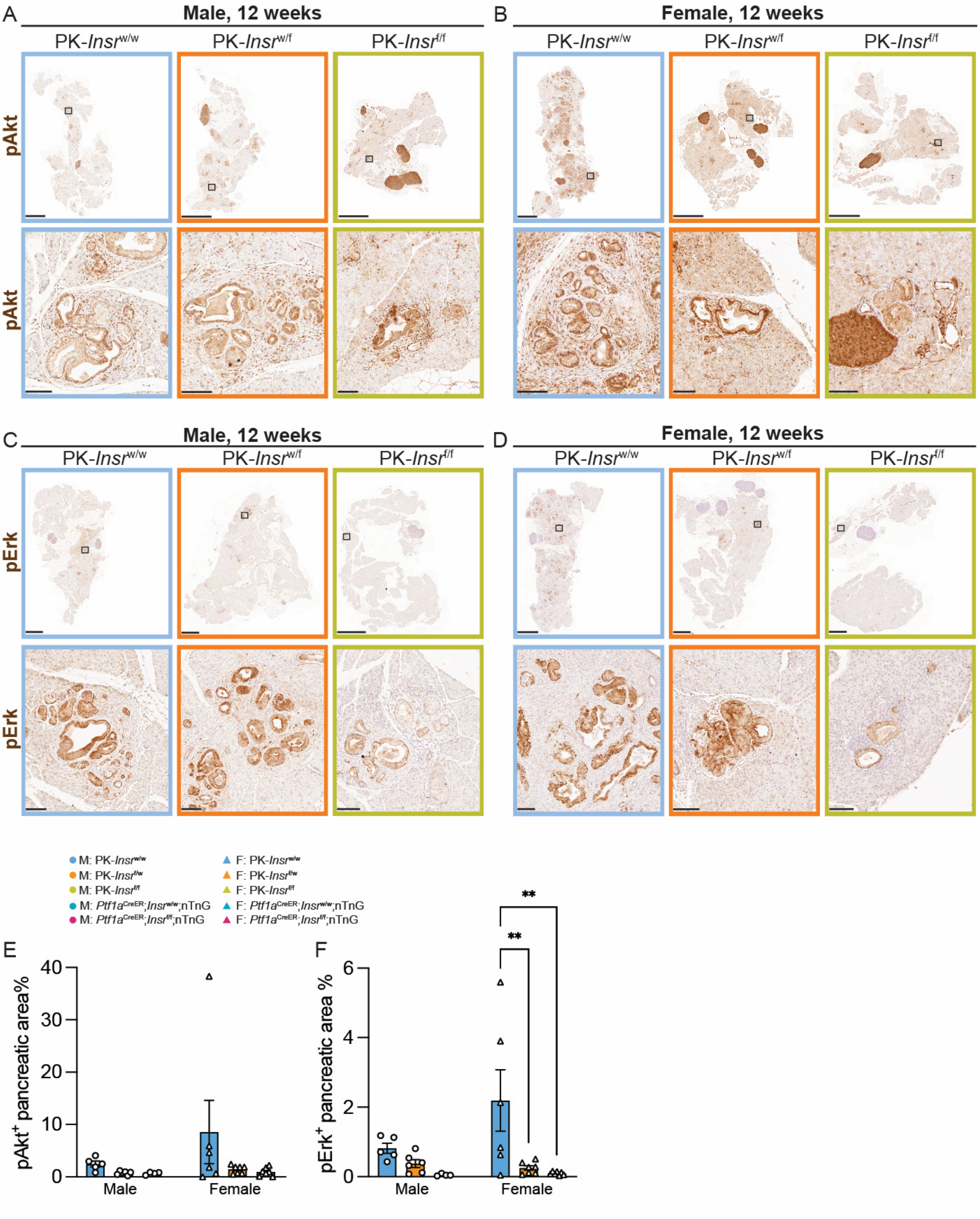

A
Male, 12 weeks
PK-Insr w/w
PK-Insr w/f
PK-Insr f/f
pAkt
B
Female, 12 weeks
C
pErk
D
M: PK-Insr w/w
M: PK-Insr f/w
M: PK-Insr f/f
M: Ptf1a CreER;Insr w/w;nTnG
M: Ptf1a CreER;Insr f/f;nTnG
F: PK-Insr w/w
F: PK-Insr f/w
F: PK-Insr f/f
F: Ptf1a CreER;Insr w/w;nTnG
F: Ptf1a CreER;Insr f/f;nTnG
E
pAkt+ pancreatic area%
0
10
20
30
40
Male
Female
F
pErk+ pancreatic area %
0
2
4
6
**
**

**Figure S4. Loss of *Insr* in acinar cells reduced pAkt and pErk, related to Figure 3.**

**A-D,** Representative whole section (top) and high-magnification (bottom) images of immunohistochemical staining for pAkt (**A-B**) or pErk (**C-D**) in 12-week-old male (**A, C**) and female (**B, D**) PK-$Insr^{w/w}$, PK-$Insr^{w/f}$, and PK-$Insr^{f/f}$ pancreata. **E-F**, Quantification of $pAkt^{+}$ (**E**) or $pErk^{+}$ (**F**) pancreatic area for mice from each genotype and sex (M or F) (n= 4-7). Scale bars: 2 mm (top) and 0.1 mm (bottom). **p<0.01 by one-way ANOVA (F).

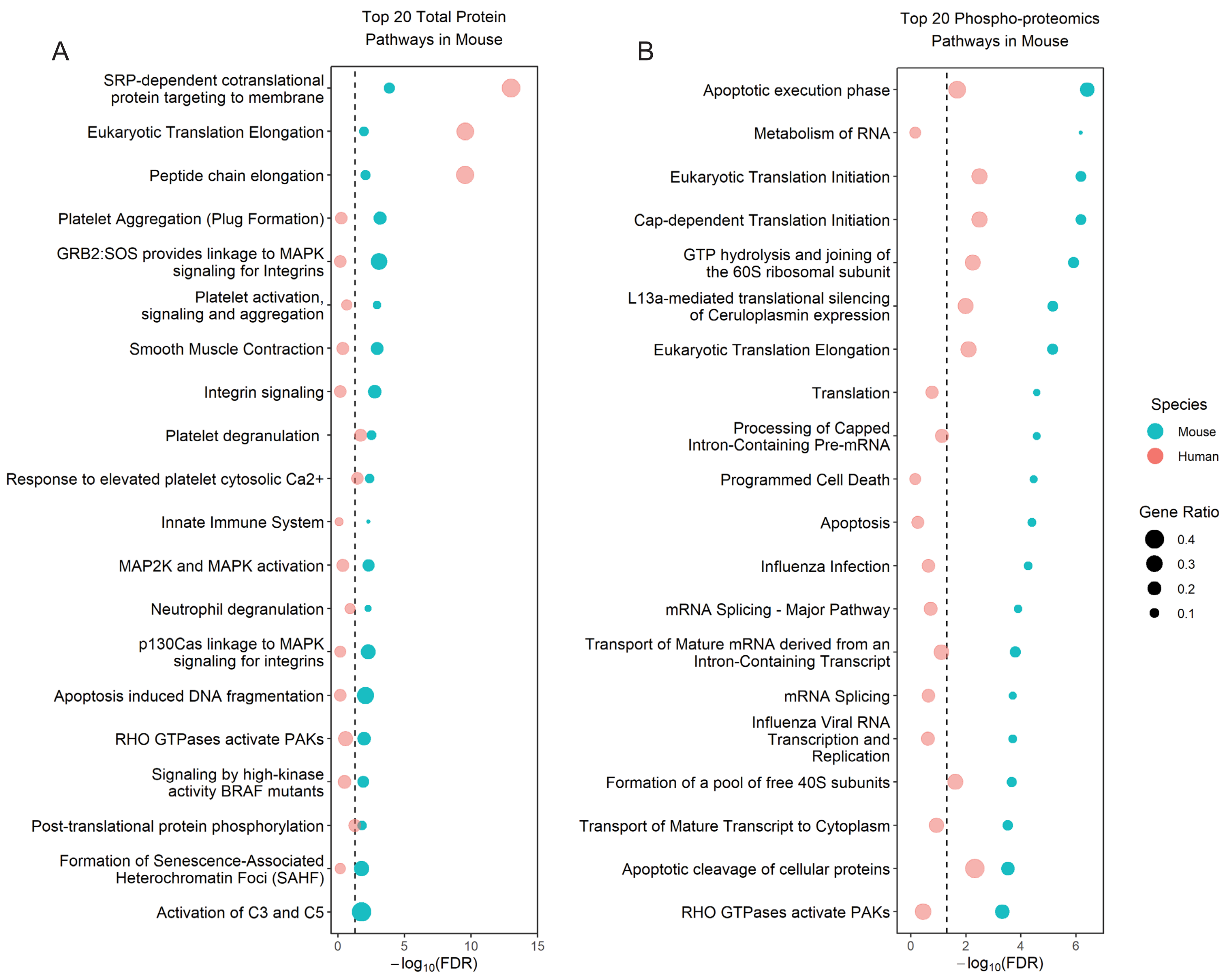

A
Top 20 Total Protein Pathways in Mouse
SRP-dependent cotranslational protein targeting to membrane
Eukaryotic Translation Elongation
Peptide chain elongation
Platelet Aggregation (Plug Formation)
GRB2:SOS provides linkage to MAPK signaling for Integrins
Platelet activation, signaling and aggregation
Smooth Muscle Contraction
Integrin signaling
Platelet degranulation
Response to elevated platelet cytosolic Ca2+
Innate Immune System
MAP2K and MAPK activation
Neutrophil degranulation
p130Cas linkage to MAPK signaling for integrins
Apoptosis induced DNA fragmentation
RHO GTPases activate PAKs
Signaling by high-kinase activity BRAF mutants
Post-translational protein phosphorylation
Formation of Senescence-Associated Heterochromatin Foci (SAHF)
Activation of C3 and C5
0
5
10
15
$-\log_{10}(FDR)$
B
Top 20 Phospho-proteomics Pathways in Mouse
Apoptotic execution phase
Metabolism of RNA
Eukaryotic Translation Initiation
Cap-dependent Translation Initiation
GTP hydrolysis and joining of the 60S ribosomal subunit
L13a-mediated translational silencing of Ceruloplasmin expression
Eukaryotic Translation Elongation
Translation
Processing of Capped Intron-Containing Pre-mRNA
Programmed Cell Death
Apoptosis
Influenza Infection
mRNA Splicing - Major Pathway
Transport of Mature mRNA derived from an Intron-Containing Transcript
mRNA Splicing
Influenza Viral RNA Transcription and Replication
Formation of a pool of free 40S subunits
Transport of Mature Transcript to Cytoplasm
Apoptotic cleavage of cellular proteins
RHO GTPases activate PAKs
0
2
4
6
$-\log_{10}(FDR)$
Species
Mouse
Human
Gene Ratio
0.4
0.3
0.2
0.1

**Figure S5.** ***Kras*$^{G12D}$ mutant mice had similar proteomics and phospho-proteomics changes as human PDAC tumor samples, related to Figure 4.**

**A,** Plot of the top 20 mouse pathways from Reactome pathway enrichment for proteins that were significantly up-or down-regulated (Adj. $p<0.05$) in PK-*Insr*$^{w/w}$ mice (n=3) compared to *Ptf1a*$^{CreER}$;*Insr*$^{w/w}$;nTnG mice (n=3). The FDR for the same pathways from pathway enrichment with proteins from human data that were significantly up- or down-regulated (Adj. $p<10^{-19}$) in normal (n=75) compared to tumor samples (n=140) are also shown. The size of each dot represents the proportion of genes inputted in all the genes found in each pathway. **B,** Plot of the top 20 mouse pathways from Reactome pathway enrichment for proteins from corresponding phospho-sites that were significantly up- or down-regulated (Adj. $p<0.05$) in PK-*Insr*$^{w/w}$ mice (n=3) compared to *Ptf1a*$^{CreER}$;*Insr*$^{w/w}$;nTnG mice (n=3). The FDR for the same pathways from pathway enrichment with proteins from corresponding phospho-sites from human data that were significantly up- or down-regulated (Adj. $p<10^{-12}$) in normal (n=75) compared to tumor samples (n=140) are also shown. The size of each dot represents the proportion of genes inputted in all the genes found in each pathway.

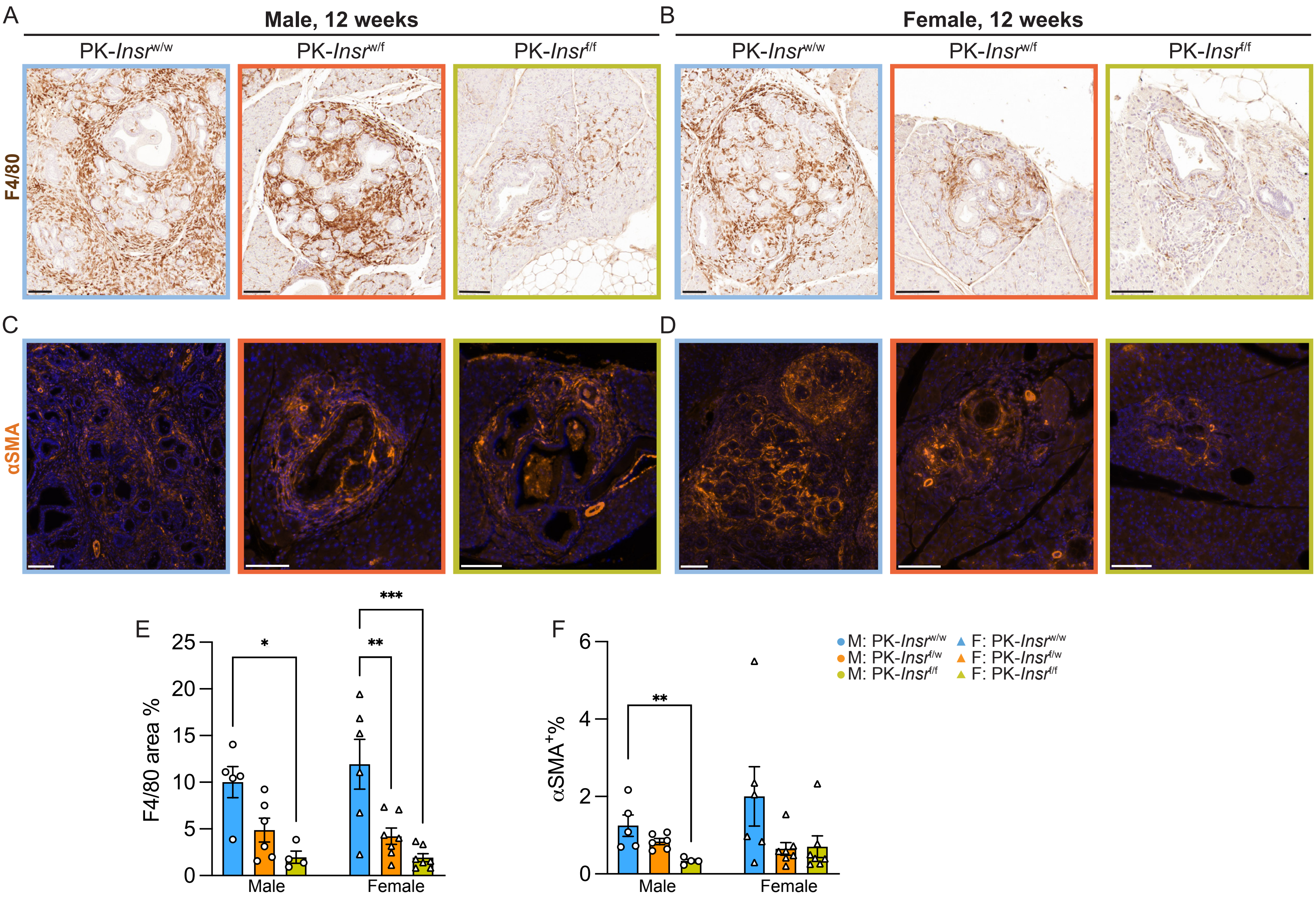

A
Male, 12 weeks
B
Female, 12 weeks
PK-Insr w/w
PK-Insr w/f
PK-Insr f/f
F4/80
C
D
αSMA
E
F4/80 area %
Male
Female
F
αSMA+%
M: PK-Insr w/w
M: PK-Insr f/w
M: PK-Insr f/f
F: PK-Insr w/w
F: PK-Insr f/w
F: PK-Insr f/f

**Figure S6. Loss of *Insr* in acinar cells reduced desmoplasia and inflammation, related to Figure 7.**

**A-B**, Representative high-magnification images of immunohistochemical staining of F4/80 (**A-B**) and $\alpha$SMA (**C-D**) for 12-week-old male (**A, C**) and female (**B, D**) PK-*Insr*$^{w/w}$, PK-*Insr*$^{w/f}$, and PK-*Insr*$^{f/f}$ mice. **E,** Quantification of F4/80$^{+}$ pancreatic area for mice from each genotype and sex (M or F) (n= 4-7). **F,** Quantification of $\alpha$SMA$^{+}$ pancreatic area for mice from each genotype and sex (M or F) (n= 4-7). *p<0.05, **p<0.01, ***p<0.001 by one-way ANOVA (E), or by Kruskal-Wallis test (F).

A

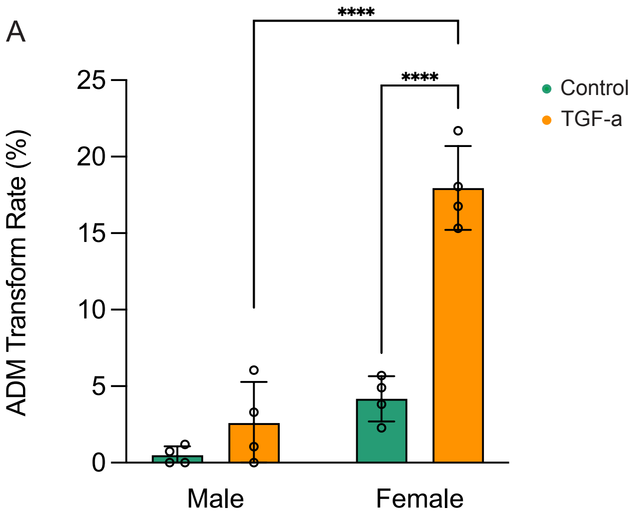

**Figure S7. Female acinar cells are more responsive to TGF-α *ex vivo* than male acinar cells, related to Figure 7.**

**A**, Quantification of ADM transformation rate of all acinar clusters per well. Acinar clusters were isolated from wild-type Bl6 female (n=4) or male (n=4) mice. Acinar cell 3D explants were quantified after 5 days treatment with a combination of ± TGF-α with soybean trypsin inhibitor in RPMI media. ****$p<0.0001$ by two-way ANOVA (A).
